# *EnvRtype*: a software to interplay enviromics and quantitative genomics in agriculture

**DOI:** 10.1101/2020.10.14.339705

**Authors:** Germano Costa-Neto, Giovanni Galli, Humberto Fanelli Carvalho, José Crossa, Roberto Fritsche-Neto

**Affiliations:** Department of Genetics, ‘Luiz de Queiroz’ Agriculture College, University of São Paulo, São Paulo, Brazil; Biometrics and Statistics Unit, Genetic Resources Program, and Global Wheat Program, International Maize and Wheat Improvement Center (CIMMYT), Texcoco, Mexico; Quantitative Genetics and Biometrics Cluster, International Rice Research Institute (IRRI), Los Baños, Philippines

**Author notes:** To whom correspondence should be addressed: Germano Costa-Neto.

**Keywords:** *G×E*: genotype × environment interaction, Envirotyping, Environmental Characterization

## Abstract

Envirotyping is an essential technique used to unfold the non-genetic drivers associated with the phenotypic adaptation of living organisms. Here we introduce the *EnvRtype* R package, a novel toolkit developed to interplay large-scale envirotyping data (enviromics) into quantitative genomics. To start a user-friendly envirotyping pipeline, this package offers: (1) remote sensing tools for collecting (get_weather and extract_GIS functions) and processing ecophysiological variables (processWTH function) from raw environmental data at single locations or worldwide; (2) environmental characterization by typing environments and profiling descriptors of environmental quality (env_typing function), in addition to gathering environmental covariables as quantitative descriptors for predictive purposes (W_matrix function); and (3) identification of environmental similarity that can be used as an enviromic-based kernel (env_typing function) in whole-genome prediction (GP), aimed at increasing ecophysiological knowledge in genomic best-unbiased predictions (GBLUP) and emulating reaction norm effects (get_kernel and kernel_model functions). We highlight literature mining concepts in fine-tuning envirotyping parameters for each plant species and target growing environments. We show that envirotyping for predictive breeding collects raw data and processes it in an eco-physiologically-smart way. Examples of its use for creating global-scale envirotyping networks and integrating reaction-norm modeling in GP are also outlined. We conclude that *EnvRtype* provides a cost-effective envirotyping pipeline capable of providing high quality enviromic data for a diverse set of genomic-based studies, especially for increasing accuracy in GP across untested growing environments.

## INTRODUCTION

Quantitative genetics divides phenotypic variation (P) into a genetic (G) and non-genetic source of variation (E). The latter may involve micro-environmental effects that can be controlled by adequate experimental designs and phenotype correction strategies (e.g., Resende and Duarte, 2007; Galli et al., 2018). Conversely, most non-genetic sources are due to macro-environmental fluctuations resulting from resource availability during crop lifetime (Shelford, 1931). Despite this unfolded division, the effect of the environment on shaping gene expression (e.g., Plessis et al., 2015; Jończyk et al., 2017; Liu et al., 2020) and fine-tuning epigenetic factors (Varotto et al., 2020; Vendramin et al., 2020) creates an indissoluble envirotype-phenotype covariance in the phenotypic records (Lynch and Walsh, 1998). Thus, for any genotype-phenotype association study across multiple environments (e.g., mapping quantitative trait loci, QLT; genomic association studies, GWAS), there is a strong non-genetic influence that can be better understood using envirotyping-based data, i.e., a foundation of multiple techniques to collect, process, and integrate environmental information in genetic and genomic studies, Costa-Neto et al. (2020a).

Over the last ten years, envirotyping (Xu, 2018) has been incorporated into whole-genome prediction (GP, Meuwissen et al., 2001) aiming to better model genotype × environment interaction (G×E) as a function of reaction-norm from environmental covariables (ECs), i.e., linearized responsiveness of a certain genotype for a target environmental gradient. Those genomic-related reaction norms can be modeled as genotype-specific coefficients for each EC due to wholegenome factorial regressions (Heslot et al., 2014; Ly et al., 2018; Millet et al., 2019), allowing a deeper understanding of which ECs may better explain the phenotypic plasticity of organisms. Furthermore, ECs can also be used to create envirotyping-based kinships (Jarquín et al., 2014; Morais-Junior et al., 2018; Costa-Neto et al., 2020a), enabling the establishment of putative environmental similarities that may drive a large amount of phenotypic variation. The integration of ecophysiological enriched envirotyping data has led to outstanding results in modeling crops such as maize, due to the use of Crop Growth Models (Cooper et al., 2016; Messina et al., 2018) and Deep Kernel approaches (Costa-Neto et al., 2020a). Combined with phenotyping and genotyping data, the use of envirotyping data may leverage molecular breeding strategies to understand historical trends and cope with future scenarios of environmental change (Gillberg et al., 2019; de los Campos et al., 2020). Its use can also support other prediction-based pipelines in plant breeding, such as high-throughput phenotyping surveys (Krause et al., 2019; Bustos-Korts et al., 2019; Galli et al., 2020).

Despite advancements in the development of hypotheses supporting the inclusion of envirotyping data in GP, it is difficult for most breeders to deal with the interplay between envirotyping, ecophysiology, and genetics. For example, much research has been conducted to explore and associate data into the concepts and theories underlying quantitative genetics (e.g., Fisher’s Infinitesimal Model) with the goal of building genomic relationship matrices (GRM). Genotyping pipelines based on bioinformatics were successfully developed to translate biochemical outputs collected from plant tissues into biologically significant markers of DNA polymorphisms, e.g., genotyping-by-sequence (GBS, Elshire et al., 2011). To the best of our knowledge, there is no publicly available user-friendly software to implement envirotyping pipelines to translate raw environmental data into a useful, highly-tailored matrix of envirotypic descriptors. Consequently, workflow to interplay enviromics (pool of environmental types, abbreviated as envirotypes) and genomic analysis is lacking, especially for GP conditions in multi-environment testing (MET) where G×E is the main hindrance in the model’s accuracy.

In this study, we introduce *EnvRtype*, a novel R package used to integrate macro-environmental factors in various fields of plant, animal, or ecological science. We approach basic ecophysiological concepts underlying the collection and processing of raw-environmental data, both biologically and statistically. Then, we present the functions for implementing remote data collection and primary processing and its applications for deriving quantitative and qualitative descriptors of relatedness. Finally, we present a comprehensive view of how envirome-based data can be incorporated into GP for selecting genotypes across diverse environments. We highlight the use of different envirotyping levels to discover descriptors of environmental similarity, using crop species to exemplify the concepts.

## MATERIAL AND METHODS

### Envirotyping Pipeline

*EnvRtype* is an R package created for handling envirotyping by ecophysiological concepts in quantitative genetics and genomics for multiple environments. It means that envirotyping is not only a collection of raw environmental data that is used for exploratory or predictive processes but rather a pipeline based on the collection of raw data and their subsequent processing in a manner that makes sense for describing the development of an organism in the target environment using *a priori* ecophysiological knowledge. Here we consider *enviromics* as the large-scale envirotyping (Xu, 2018; Resende et al., 2020) of a theoretical population of environments for a target species or germplasm (the so-called envirome). It may also denote the core of possible growing conditions and technological inputs that create different productivity levels. The envirotyping pipeline implemented by *EnvRtype* software is divided into three modules briefly described above and detailed in the following sections (Fig. 1).

**Figure 1.**
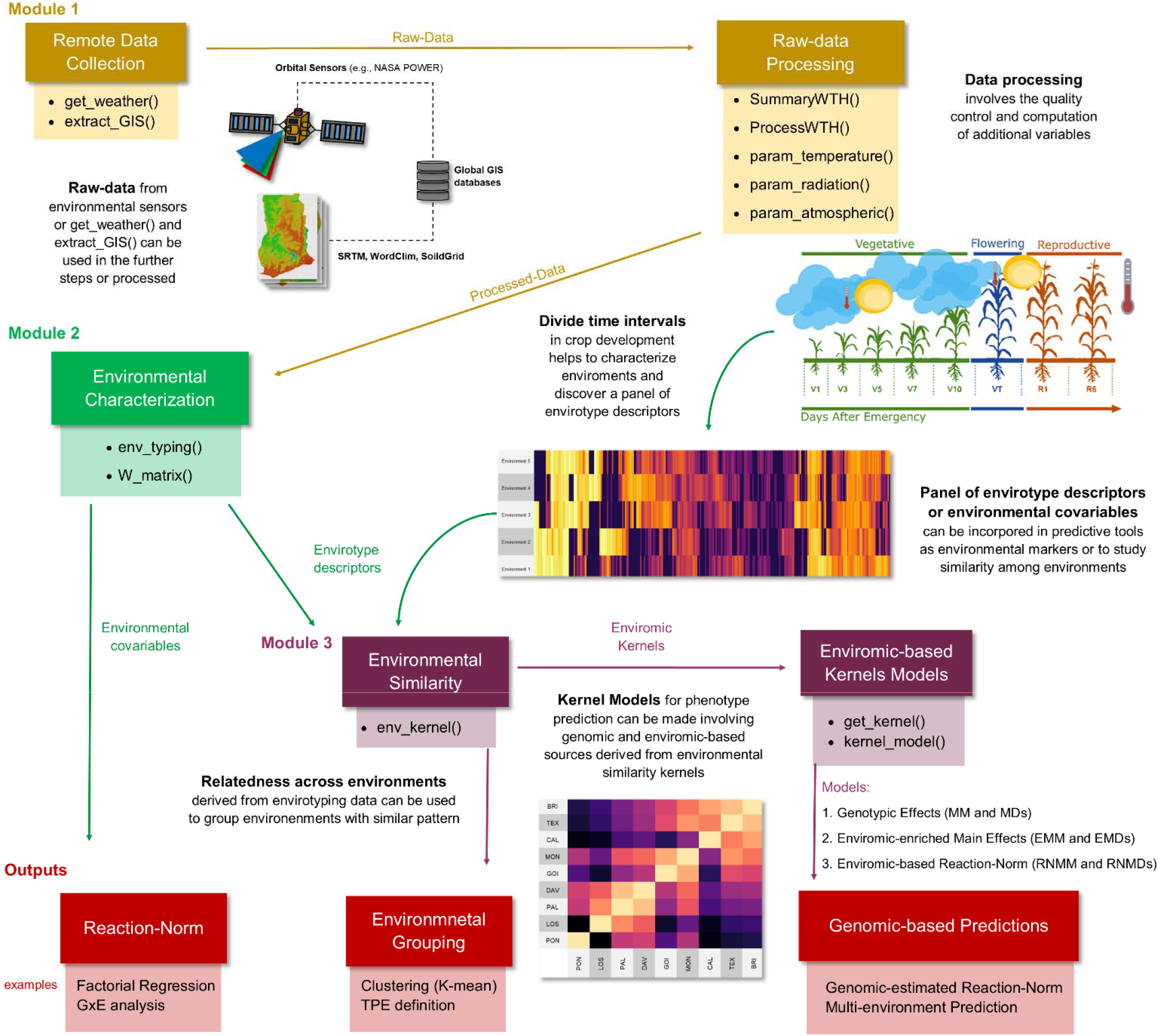
The workflow of the envirotyping pipeline implemented using *EnvRtype* in R. Yellow, green, dark purple, and red-colored boxes denote the steps related to Modules 1, 2, 3, and the examples of outputs from *EnvRtype*. Straight arrows indicate the flux of the envirotyping pipeline passing by each module. A curved arrow represents a process between Modules 1 and 2 in which field-growing conditions can be described as a panel of envirotype descriptors from each environmental factor processed and organized in Module 2.

Module 1 (yellow toolboxes in Fig. 1) starts by collecting raw environmental data from public platforms, such as a satellite-based weather system named ‘NASA’s Prediction of Worldwide Energy Resources’ (NASA POWER, https://power.larc.nasa.gov/), which can access information daily anywhere on earth. This database is well consolidated and validated for use in several research fields, including crop modeling in agricultural research (White et al., 2011; Monteiro et al., 2018; Aboelkhair et al., 2019). Details about resolution and validation are given in https://power.larc.nasa.gov/docs/methodology/validation/.

Data collection may span existing experimental trials (single sampling trials) or historical trends for a given location × planting date arrangement. This module gathers the functions for remote data collection of daily weather and elevation data, as well as the computation of ecophysiological variables, such as the effect of air temperature on radiation use efficiency. The module includes a toolbox with ‘Remote Data Collection’ and ‘Data Processing’ steps, both designed to help researchers find a viable alternative for expensive in-field environmental sensing equipment. More details about the theoretical basis of environmental sensing and the module are given in the section **Module 1: Remote Environmental Sensing**.

The processed environmental information can then be used for many purposes. In Module 2, we designed tools for the characterization of the macro-environmental variations, which can also be done across different time intervals of crop growth and development (when associated with a crop) or fixed time intervals (to characterize locations). The environmental characterization toolbox (green toolbox in Fig. 1) involves two types of profiling:

1. Discovering environmental types (envirotypes, hereafter abbreviated as ETs) and how frequently they occur at each growing environment (location, planting date, year). Based on the ET-discovering step, it is possible to create environmental profiles and group environments with the same ET pattern. This step is also useful for running exploratory analysis, e.g., to discover the main ET of planting dates at a target location.
2. Gathering environmental covariables (hereafter abbreviated as ECs) from point-estimates (e.g., mean air temperature, cumulative rainfall). These ECs can be used for many purposes, from a basic interpretation of G×E to estimating gene-environment interactions. At the end of this process, a matrix of ECs (**W**) is created and integrated with tools from Module 3.

Further details about this module are given in the section **Module 2: Macro-Environmental Characterization**.

Finally, the information from Module 2 can be used to create environmental similarity and integrate robust GP platforms for multiple environments, hereafter referred to as envirotype-informed GPs. Module 3 (the dark purple boxes Figure 1) aims to provide tools to compute environmental similarity using correlations or Euclidean distances across different trials conducted on ECs. Thus, we developed a function to integrate this enviromic source in GP as an additional source of variation to bridge the gap between genomic and phenotypic variation. For that, we provide at least four different structures into a flexible platform to integrate multiple genomic and enviromic kinships.

Figure 1 shows some possible outputs of the *EnvRtype* package (in red toolbox colors), in which **W** can be used to interpret G×E (e.g., factorial regression) or exploit it in terms of increasing the accuracy of phenotype prediction for multiple environments. More details are given in the section **MODULE 3: Enviromic Similarity and Phenotype Prediction.** Below we give some theoretical details about each module and a description of the functions used to implement it.

### Software

The R package *EnvRtype* is available at https://github.com/allogamous/EnvRtype [version 0.1.6, verified December 15th, 2020]). More details about graphical plots and additional codes can also be found on this Git Hub webpage and in Supplementary Software. Typing the following command in R will automatically install the package (**BOX 1**):

#### BOX 1: Install EnvRtype

~~~
> install.packages(‘devtools’); devtools::install_github(‘allogamous/EnvRtype’)
~~~

All codes and BOX in this work are available on both the Git Hub web page and as Supplementary Code.

### Data sets and Codes availability

*EnvRtype* has a toy data set for running examples, mostly involving genomic prediction (see Module 3). This data set was included in Souza et al. (2017) and Cuevas et al. (2019) and came from the Helix Seed Company (HEL). However, to facilitate the demonstration of functions, we provided a subset of 150 hybrids per environment (**BOX 2**). Grain yield data are mean-centered and scaled (*MaizeYield* object). The genotyping relationship for additive effects is based on 52,811 SNPs available to make the predictions (*maizeG* object). The phenotypic and genomic data are credited to Helix Seed Ltda. Company. Finally, weather data are presented for each of the five environments (*maizeWTH* object). All codes are available in the BOX codes (from 1 to 19), and as Supplementary Codes. Additional tutorials can be found at Git Hub (https://github.com/allogamous/EnvRtype).

#### BOX 2: Data sets

~~~
> data(‘maizeYield’) # toy set of phenotype data (grain yield per environment)
> data(‘maizeG’) # toy set of genomic relationship for additive effects
> data(‘maizeWTH’) # toy set of environmental data
~~~

### MODULE 1: Remote Environmental Sensing

#### Remote data collection

*EnvRtype* implements the remote collection of daily weather and elevation data by the *get_weather* function. This function has the following arguments: the environment name (**env.id**); geographic coordinates (latitude, **lat**; longitude, **lon**) in WGS84; time interval (**start.day** and **end.day**, given in ‘year-month-day’); and country identification (**country**), which sets the raster file of elevation for the region of a specific country. Countries are specified by their 3-letter ISO codes (check in the package Git Hub or use the getData(‘ISO3’) function from the raster package to see the codes).

Table 1 shows the names of the outputs of *get_weather* and *processWTH* (see Tools for basic processing). All weather information is given on a daily basis. Altitude (*ALT*) information is provided by SRTM 90 m resolution and can be collected from any place between −60 and 60 latitudes. This information is presented as a *data.frame* class output in R. It is possible to download data for several environments by country.

**Table 1.** The core of environmental factors available using the ‘Environmental Sensing Module’ of the *EnvRtype* package.

| Source | Environmental Factor | Unit |
| --- | --- | --- |
| Nasa<br>POWER<sup>1</sup> | Top-of-atmosphere insolation | MJ m<sup>-2</sup> d<sup>-1</sup> |
|  | Average insolation incident on a horizontal surface | MJ m<sup>-2</sup> d<sup>-1</sup> |
|  | Average downward longwave radiative flux | MJ m<sup>-2</sup> d<sup>-1</sup> |
|  | Wind speed at 10 m above the surface of the earth | m s<sup>-1</sup> |
|  | Minimum air temperature at 2 m above the surface of the earth | °C d<sup>-1</sup> |
|  | Maximum air temperature at 2 m above the surface of the earth | °C d<sup>-1</sup> |
|  | The dew-point temperature at 2 m above the surface of the earth | °C d<sup>-1</sup> |
|  | Relative air humidity at 2 m above the surface of the earth | % |
|  | Rainfall precipitation (P) | mm d<sup>-1</sup> |
| SRTM<sup>2</sup> | Elevation (above sea level) | m |
| Computed<sup>3</sup> | Effect of Temperature on Radiation use Efficiency (FRUE) | - |
|  | Evapotranspiration (ETP) | mm d<sup>-1</sup> |
|  | Atmospheric water deficit P-ETP | mm d<sup>-1</sup> |
|  | The deficit of vapor Pressure | kPa d<sup>-1</sup> |
|  | The slope of saturation vapor pressure curve | kPa C° d<sup>-1</sup> |
|  | Temperature Range | °C d<sup>-1</sup> |
|  | Global Solar Radiation based on Latitude and Julian Day | MJ m<sup>-2</sup> d<sup>-1</sup> |

<sup>1</sup> collected from NASA orbital sensors (Sparks, 2018); <sup>2</sup> Shuttle Radar Topography Mission integrated with the raster R package; <sup>3</sup> processed using concepts from Allen et al. (1998) and Soltani and Sinclair (2012).

A practical example of get_weather is given below (**BOX 3**). A collection of environmental data for Nairobi, Kenya (latitude 1.367 N, longitude 36.834 E) from 01 march 2015 to 01 April 2015, is performed by:

#### BOX 3: Practical use of get_weather

~~~
> env.data = get_weather(env.id = ‘NAIROBI’,lat = −1.367,lon = 36.834,start.day = ‘2015-03-01, end.day = ‘2015-04-01’, country = ‘KEN’)
~~~

A second function is *extract_GIS*, which can collect point values from large raster files from GIS databases. This function has six arguments. The object **env.data** indicates the name of the environmental dataset (arranged as a *data.frame*). It can be an output data.frame of the *get_weather* function or any spreadsheet of environmental data, as long as it is organized with a column denoting the environment’s name, which is defined by the **env.id** argument (default is env.id = ‘env’). **Latitude** and **Longitude** is provided in the same manner described in *get_weather*.

Finally, the **name.out** is the argument to define the name of the collected covariable (e.g., ALT for altitude). The function *extract_GIS* can be useful for collecting covariables from raster files within databases such as WorldClim (Fick et al., 2017; https://www.worldclim.org/), SoilGrids (https://soilgrids.org/), EarthMaps (https://earthmap.org/) and Nasa Power (https://power.larc.nasa.gov).

A practical use of *extract_GIS* is given below (**BOX 4**). A collection of clay content (g/kg) from 5cm to 15cm of depth for Nairobi using a raster file was downloaded from *SoilGrids* and the function *extract_GIS*. The raster file ‘clay_5_15’ can be accessed in R by typing data(“clay_5_15”).

#### BOX 4: Practical use of extract_GIS

~~~
> data(“clay_5_15”)
> env.data = extract_GIS(covraster = clay_5_15c,name.out = ‘clay_5_15’,env.data = env.data)
> head(env.data)
~~~

#### Summarizing raw-data

A basic data summary of the outputs from the *get_weather* function is done by the *summaryWTH* function. This function has 10 arguments (**env.data**, **id.names**, **env.id**, **days.id**, **var.id**, **statistic, probs, by.interval, time.window,** and **names.window**). The common arguments with *extract_GIS* have the same described utility. Other identification columns (year, location, management, responsible researcher, etc.) may be indicated in the **id.names** argument, e.g., id.names = c(‘year’,’location’,’treatment’).

Considering a specific environmental variable, the argument **var.id** can be used as, for example, var.id = ‘T2M’. By default, this function considers all the names of the variables presented in Table 1. For other data sources, such as micro-station outputs, this argument is necessary for identifying which variables will be summarized. The argument **days.id** indicates the variable pointing to the time (days), where the default is the *daysFromStart* column from the *get_weather* function. A basic example of this use is given below (**BOX 4**):

#### BOX 4: Practical use of SummaryWTH

~~~
> summaryWTH(env.data = env.data, env.id = ‘env’, days.id = ‘daysFromStart’,statistic = ‘mean’)
> summaryWTH(env.data = env.data) # by default
~~~

Dividing the development cycle into time intervals (e.g., phenology), whether phenological or fixed time intervals (e.g., 10-day intervals), helps to understand the temporal variation of environmental factors during the crop growth cycle. Thus, specific time intervals can be created by the **time.window** argument. For example, time.window = c(0,14,35,60,90,120) denotes intervals of 0-14 days from the first day on record (0). If the first record denotes the crop’s emergence date in the field, this can also be associated with some phenological interval. Those intervals can be named using the argument **names.window**, names.window = c(‘P-E’,’E-V1’,’V1-V4’,’V4-VT’,’VT-GF’,’GF-PM’).

The argument **statistic** denotes which statistic should be used to summarize the data. The statistic can be: *mean, sum* or *quantile*. By default, all statistics are used. If statistic = ‘quantile’, the argument **prob** is useful to indicate which percentiles (from 0 to 1) will be collected from the data distribution, i.e., default is prob = c(0.25, 0.50, 0.75), denoting the first (25%) second (50%, median) and third (75%) quantiles.

#### Tools for basic data processing

The *processWTH* function performs basic data *process*ing. As described for *summaryWTH*, this function can also process environmental data for get_weather outputs and other sources (micro-stations, in-field sensors) using the same identification arguments (**env.data**, **id.names**, **env.id**, **days.id**, **var.id**). This function also gathers three other sub-functions created to compute general variables related to ecophysiological processes, such as the macro effects of soil-plant-atmosphere dynamics and atmospheric temperature on crop development. The basic usage of this package is given by processWTH(env.data = env.data); notwithstanding, crop-specific parameters such as cardinal values of temperature and evapotranspiration, as well as site-specific characteristics, can be given in additional arguments. Below, we describe these arguments over three functions that compose processWTH. We provide a brief description about them, in addition to the ecophysiological concepts underlying their application (Appendix).

#### Radiation-related covariables

EnvRtype made a function called *param_radiation* available to compute additional radiation-based variables that can be useful for plant breeders and researchers in several fields of agricultural research (e.g., agrometeorology). These parameters include the actual duration of sunshine hours (*n*, in hours) and total daylength (*N*, in hours), both estimated according to the altitude and latitude of the site, time of year (Julian day, from 1 to 365), and cloudiness (for *n*). In addition, the global solar radiation incidence (SRAD, in MJ m^2^ d^-1^) is computed as described at the beginning of this section. The latter is important in most computations of crop evapotranspiration (Allen et al., 1998) and biomass production (Muchow et al., 1990; Muchow and Sinclair, 1991). More details about those equations are given in ecophysiology and evapotranspiration literature (Allen et al., 1998; Soltani and Sinclair, 2012).

The arguments of *param_radiation* are: **env.data** and **merge**, in which merge denotes if the computed radiation parameters must be merged with the env.data set (merge = TRUE, by default).

#### Temperature-related covariables

The *param_temperature* function computes additional thermal-related parameters, such as *GDD* and *FRUE*, and *T2M_RANGE*. This function has eight arguments (**env.data**, **Tmax, Tmin**, **Tbase1**, **Tbase2**, **Topt1**, **Topt2** and **merge**). To run this function with data sources other than get_weather, it is necessary to indicate which columns denote maximum air temperature (Tmax, default is Tmax = ‘T2M_MAX’) and minimum air temperature (Tmin, default is Tmin = ‘T2M_MIN’) (**BOX 6**). The cardinal temperatures must follow the processes provided in the previously described ecophysiology literature (Soltani and Sinclar, 2012, see Appendix 3). Consider the estimations for dry beans at the same location in Nairobi, Kenya (previous box examples). The cardinal temperatures for dry beans were collected from Table 2, which gathers different cardinal temperatures for a diverse set of crops.

#### BOX 6: Practical use of param_temperature for Dry Beans in Nairobi, Kenya

~~~
> TempData = param_temperature(env.data = env.data,Tbase1 = 8,Tbase2 = 45,Topt1 = 30,Topt2 = 35)
> head(TempData)
> env.data = param_temperature(env.data = env.data,Tbase1 = 8,Tbase2 = 45,Topt1 = 30,Topt2 = 35,merge = TRUE) # merging TempData automatically
> head(env.data)
~~~

**Table 2.** Synthesis of some cardinal limits for temperature on the phenology development in the main crops. These estimates were adapted from Soltani and Sinclar (2012), Lago et al. (2009), and Bartz et al. (2017).

| Specie | Suggested Cardinal Limit |  |  |  |
| --- | --- | --- | --- | --- |
|  | Tbase1 | Topt1 | Topt2 | Tbase2 |
| Maize | 8.0 | 30.0 | 37.0 | 45.0 |
| Wheat | 0.0 | 25.0 | 28.0 | 40.0 |
| Rainfed Rice | 8.0 | 30.0 | 37.0 | 45.0 |
| Irrigated Rice (only vegetative stage) | 8.0 | 28.0 | 40.0 | 45.0 |
| Irrigated Rice (only reproductive stage) | 15.0 | 25.0 | 35.0 | 45.0 |
| Sorghum | 8.0 | 30.0 | 37.0 | 45.0 |
| Soybean | 8.0 | 30.0 | 35.0 | 45.0 |
| Peanut | 8.0 | 30.0 | 35.0 | 45.0 |
| Canola | 0.0 | 25.0 | 28.0 | 40.0 |
| Sunflower | 8.0 | 30.0 | 34.0 | 45.0 |
| Dry Bean | 8.0 | 30.0 | 35.0 | 45.0 |
| Chickpea | 0.0 | 25.0 | 30.0 | 40.0 |
| Barley | 0.0 | 25.0 | 28.0 | 40.0 |
| Sugarcane | 5.0 | 22.5 | 35.0 | 40.0 |

#### Atmospheric demands

We implemented the *param_atmospheric* function to run basic computations of atmospheric demands. This function has 11 arguments: **env.data**; **PREC** (rainfall precipitation in mm, default is PREC=‘PRECTOT’); **Tdew** (dew point temperature in °C, default is Tdew=‘T2M_DEW’); **Tmax** (maximum air temperature in °C, default is Tmax=‘T2M_MAX’); **Tmin** (minimum air temperature in °C, default is Tmin=‘T2M_MIN’); **RH** (relative air humidity %, default is RH=‘RH2M’); **Rad** (net radiation, in MJ m^-2^ day^-1^, default is Rad =‘Srad’); **alpha** (empirical constant accounting for vapor deficit and canopy resistance values, default is alpha=1.26); **Alt** (altitude, in meters above sea level, default is Alt = ALT); **G** (soil heat flux in W m^-2^, default is G=0); and **merge** (default is merge=TRUE). Details about these inputs and equations are given in the Appendix Section.

Below we present an example of usage in Nairobi, Kenya. Consider the same env.data collected in the previous box and an elevation value of Alt = 1,628 (**BOX 7**).

#### BOX 7: Practical use of param_atmospheric for Dry Bean Crop in Nairobi, Kenya

~~~
> RadData = param_radiation(env.data = env.data) # first need to compute radiation parameters
> head(RadData)
> env.data = param_radiation(env.data = env.data,merge = TRUE) # need first some radiation inputs
> AtmData = param_atmospheric(env.data = env.data, Alt = 1628)
> head (AtmData)
> env.data = param_atmospheric(env.data = env.data, Alt = 1628,merge = TRUE)
> head(env.data)
~~~

### MODULE 2: Macro-Environmental Characterization

#### Discovering Envirotypes with env_typing

An environment can be viewed as the status of multiple resource inputs (e.g., water, radiation, nutrients) across a certain time interval (e.g., from sowing to harvesting) within a specific space or location. The quality of those environments is an end-result of the daily balance of resource availability, which can be described as a function of how many resources are available and how frequently those resources occur (e.g., transitory or constant effects). Also, the relationship between resource absorption and allocation depends on plant characteristics (e.g., phenology, current health status). Then, this particular environmental-plant influence is named after the envirotype to differentiate it from the concept of raw environmental data (data collected directly from sensors). It can be referred to as environmental type (ET). Finally, the typing of environments can be done by discovering ETs; the similarity among environments is a consequence of the number of ETs shared between environments.

Before the computation of ETs, a first step was to develop an design based on ecophysiological concepts (e.g., plants’ needs for some resource) or summarize the raw data from the core environments being analyzed. Then, for each ET we computed the frequency of occurrence, which represents the frequency of specific quantities of resources available for plant development. Typing by frequency of occurrence provides a deeper understanding of the distribution of events, such as rainfall distribution across different growing cycles and the occurrence of heat stress conditions in a target location (Heinemann et al. 2015). Thus, groups of environments can be better identified by analyzing the events occurring in a target location, year, or planting date. This step can be done not only by using grade point averages (e.g., accumulated sums or means for specific periods), but also by their historical similarity. In this way, we are able to not only group environments in the same year, but also through a historical series of years. Finally, this analysis deepens in resolution when the same environment is divided by time intervals, which can be fixed (e.g., 10-day intervals), or categorized by specific phenological stages of a specific crop.

To implement envirotype profiling, we created the *env_typing* function. This function computes the frequency of occurrence of each envirotype across diverse environments. This function has 12 arguments, nine of which (**env.data**, **id.names**, **env.id**, **days.id var.id**, **statistic, by.interval, time.window, and names.window**) work in the same way as already described in the previous functions. The argument **cardinals** are responsible for defining the biological thresholds between envirotypes and adaptation zones. These cardinals must respect the ecophysiological limits of each crop, germplasm, or region. For that, we suggest literature on ecophysiology and crop growth modeling, such as Soltani and Sinclar (2012). The argument **cardinals** can be filled out as vectors (for single-environmental factors) or as a list of vectors for each environmental factor considered in the analysis. For example, considering the cardinals for air temperature in rainfed rice presented in Table 2, the cardinals are typed for Los Baños, Philippines, from 2000 to 2020, as (**BOX 8**):

#### BOX 8: Basic use of env_typing for typing temperature in Los Baños, Philipines, from 2000 to 2020

~~~
> env.data = get_weather(env.id = ‘LOSBANOS’,country = ‘PHL’,
>   lat = 14.170,lon = 121.241,variables.names = ‘T2M’,
>   start.day = ‘2000-03-01’,end.day = ‘2020-03-01’)
> card = list(T2M=c(0,8,15,28,40,45,Inf)) # a list of vectors containing empirical and cardinal thresholds
> env_typing(env.data = env.data,env.id = ‘env’, var.id = ‘T2M’, cardinals = card)
~~~

If cardinals = NULL, the quantiles 10%, 25%, 50%, 75% and 90% are used by default. Which quantiles will be used is determined in the same manner as prob (in *summaryWTH*), but now using the **quantile** argument, e.g., quantile = c(0.25,0.50,0.75).

For multiple environmental factors, a list of cardinals must be provided—for example, considering rainfall precipitation (*PRECTOT*, mm.day^-1^) and dew point temperature (*T2DEW*, °C.day^-1^). Suppose precipitation values less than 10 mm.day^-1^ are insufficient to meet the studied crops’ demand. Values between 11 mm.day^-1^ and 40 mm.day^-1^ would be considered excellent water conditions, and values greater than 40 mm.day^-1^ would be considered excessive rainfall. In this scenario, such rainfall values could be negatively associated with flooding of the soil and drainage of fertilizers, among other factors related to crop lodging or disease occurrence. Thus, for PRECTOT, the cardinals will be cardinals = c(0,5,10,25,40,100). For dew point, let’s assume data-driven typing (cardinals = NULL) using the previously described quantiles. Taking the same example for Los Baños, Philippines (**BOX 9**).

#### BOX 9: Basic use of env_typing for more than one variable

~~~
> var = c(“PRECTOT”, “T2MDEW”) # variables
> env.data = get_weather(env.id = ‘LOSBANOS’,country = ‘PHL’,
>   lat = 14.170,lon = 121.241,variables.names = var,
>   start.day = ‘2000-03-01’,end.day = ‘2020-03-01’)
>card = list(PRECTOT = c(0,5,10,25,40,100), T2MDEW = NULL) # cardinals and data-driven limits
>env_typing(env.data = env.data,env.id = ‘env’, var.id = var, cardinals = card)
~~~

#### Environmental Covariables with W_matrix

The quality of an environment is measured by the amount of resources available to fulfill the plants’ demands. In an experimental network composed of multi-environment trials (MET), the environment’s quality is relative to the global environmental gradient. Finlay and Wilkinson (1963) proposed using phenotypic data as a quality index over an implicit environmental gradient. However, this implicit environmental quality index was proposed as an alternative to explicit environmental factors, given the difficulties in obtaining high-quality envirotyping data. Here we provide the use of detailed environmental data arranged in a quantitative descriptor such as a covariate matrix (**W**), following the terminology used by Costa-Neto et al. (2020a) and de los Campos et al. (2020). Based on these **W** matrices, several analyses can be performed, such as (1) dissecting the G×E interaction; (2) modeling genotype-specific sensibility to critical environmental factors; (3) dissecting the environmental factors of QTL×E interaction; (4) integrating environmental data to model the gene × environment reaction-norm; (5) providing a basic summary of the environmental gradient in an experimental network; (6) producing environmental relationship matrices for genomic prediction.

To implement these applications, the processed environmental data must be translated into quantitative descriptors by summarizing cumulative means, sums, or quantiles, such as in *summaryWTH*. However, these data must be mean-centered and scaled to assume a normal distribution and avoid variations due to differences in scale dimensions. To create environmental similarity kernels, Costa-Neto et al. (2020a) suggested using quantile statistics to better describe each variable’s distribution across the experimental network. Thus, this allows a statistical approximation of the environmental variables’ ecophysiological importance during crop growth and development. In this context, we developed the *W_matrix* function to create a double-entry table (environments/sites/years environmental factors). Conversely to *env_typing*, the *W_matrix* function was designed to sample each environmental factor’s quantitative values across different environments.

The same arguments for the functions *summaryWTH* and *env_typing* are applicable (**env.data**, **id.names**, **env.id**, **days.id var.id**, **statistic, by.interval, time.window, and names.window**). However, in *W_matrix*, arguments **center** = TRUE (by default) and **scale** = TRUE (by default) denote mean-centered 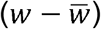 and scaled 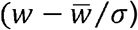, in which *w* is the original variable, 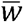 and *σ* are the mean and standard deviation of this covariable across the environments. Quality control (**QC** = TRUE argument) is done by removing covariables with more than □*σ_TOL_*±□ *σ*, where *σ_TOL_* is the tolerance limit for standard deviation, settled by default argument as *sd.tol* = 3.

To exemplify a basic use of *W_matrix*, let us consider the *maizeWTH* object, involving only weather variables temperature, rainfall and precipitation, while assuming a quality control of *sd.tol* = 4. The time intervals were settled for every ten days (default), and statistic as ‘mean’ for each variable at each time interval (**BOX 10**).

#### BOX 10: Basic use of env_typing for more than one variable

~~~
> data(“maizeWTH”) # toy set of environmental data
> var = c(“PRECTOT”, “T2MDEW”, “T2M_MAX”, “T2M_MIN”) # variables
> W = W_matrix(env.data = maizeWTH[maizeWTH$daysFromStart < 100,],
>   var.id=var, statistic=“mean”, by.interval=TRUE)
> dim(W)
~~~

### MODULE 3: Enviromic Similarity and Phenotype Prediction

The prediction of phenotypes across multiple environments can be conducted using different approaches, such as mechanistic crop models and empirical regressions, in which environmental and/or genomic information is necessary for training accurate models. The latter, named after whole-genome prediction (GP, Meuwissen et al., 2001), has revolutionized both plant and animal breeding pipelines around the world. Most approaches rely on increasing the accuracy of modeling genotype-phenotype patterns and exploring them as a predictive breeding tool. Among the several enrichments of computational efficacy and breeding applications, the integration of genomic by environment interaction (G×E) has boosted the ability of genomic-assisted selection to evaluate a wide number of genotypes under several growing conditions over multiple environmental trials (MET).

Heslot et al. (2014) and Jarquín et al. (2014) introduced environmental covariables to model an environmental source of the phenotypic correlation across MET. These approaches aim to model the reaction-norm of genotypes across MET, i.e., how different genotypes react to different environmental gradient variations. In most cases, reaction-norm modeling serves as an additional source of variation for complementing the genomic relatedness among individuals tested and untested under known environmental conditions. Thus, in addition to the genomic kernels, envirotype-informed kernels can be used to capture macro-environmental relatedness that shapes the phenotypic variation of relatives, the so-called *enviromic kernel* (Costa-Neto et al., 2020a).

In the third module of the *EnvRtype* package, we present the tools used to implement this modeling approach. Three main functions were designed for this purpose. First, the function *env_kernel* can be used for the construction of environmental relationship kernels using environmental information. Second, the *get_kernel* aims to integrate these kernels into statistical models accounting for different structures capable of explaining the phenotypic variation across MET. Finally, the function *kernel_model* can be used to fit regression models accounting for environmental and or genomic data using a computationally efficient Bayesian approach. In the following subsections, we describe the kernel methods for modeling envirotype relatedness. Then we present the statistical models that can be built with these kernels.

#### Enviromic Kernels with env_kernel

In this package, we use two types of kernel methods to compute enviromic-based similarity. The first consists of the traditional method based on the linear variance-covariance matrix (Jarquín et al., 2014). This kernel is equivalent to a genomic relationship matrix and can be described mathematically as:

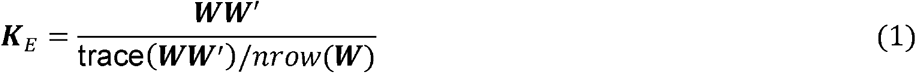

where ***K**_E_* is the enviromic-based kernel for similarity among environments and ***W*** matrix of ECs. Note that we use ***W*** matrix, but any other source of data from environments can be used here as EC (e.g., typologies, disease evaluations, management).

The second method is a nonlinear kernel modeled by Gaussian processes, commonly called the Gaussian Kernel or GK, and widely used in genomic-enabled prediction (Gianola and van Kaam (2008); de los Campos et al., 2010; Cuevas et al., 2017). The use of GK for modeling ***K**_E_* was proposed by Costa-Neto et al (2020a) and is described in a similar way to the approach already used for modeling genomic effects:

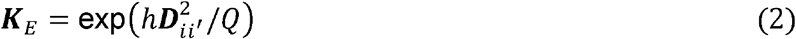

where *h* is the bandwidth factor (assume as *h* = 1 by default) factor multiplied by the Euclidean Distance 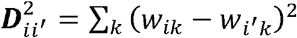 for each pairwise elements in the ***W*** = {*w_i_, w_i′_*}. This means that the environmental similarity is a function of the distance between environments realized by ECs. The scalar variable *Q* denotes the quantile used to ponder the environmental distance (assumed as *Q* = 0.5, equal to the median value of 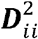. The h can be computed using a marginal function described by Pérez-Elizalde et al. (2015).

The *env_kernel* function implements both methods. It has the following main arguments: **env.data**, **env.id**, **gaussian,** and **h.gaussian**. The first two arguments work in the same manner previously described for other functions. The gaussian argument (default is gaussian = FALSE) denotes if models (1) or (2) are used to compute ***K**_E_*. If gaussian = TRUE, then the Gaussian kernel (equation 2) is used, and h.gaussian must be inserted to compute it. In the **Y** argument (default is Y = NULL), it is possible to insert a phenotypic record to be used in the marginal function to compute a data-driven *h* (Pérez-Elizalde et al., 2015).

The *env_kernel* function has two outputs called *varCov* (relatedness among covariables) and *envCov* (relatedness among environments). The first is useful to deepen the understanding of the relatedness and redundancy of the ECs. The second output is ***K_E_***. This matrix is the enviromic similarity kernel integrated into the GP models.

A basic use of env_kernel is presented below. Consider the **W** matrix created in **BOX 10** for the *maizeWTH* object (5 environments in Brazil). The ***K_E_***. value using linear covariance and the Gaussian kernel is given as (**BOX 11**):

#### BOX 11: Basic use of env_kernel for linear and nonlinear kernel methods

~~~
> env_kernel(env.data = W, gaussian = FALSE)
> env_kernel(env.data = W, gaussian = TRUE)
~~~

#### Phenotype prediction across multiple environments

After constructing the relationship kernels for environmental relatedness, it is possible to fit a vast number of statistical models using several packages available in R CRAN. However, it is important to consider that statistical models containing more complex structures (e.g., more than one genetic effect plus G×E and environmental information) are models that require more expensive computational effort and time. Under Bayesian inference, which demands multiple iterative sampling processes (e.g., via Gibbs sampler) to estimate the variance components, the computational effort may be more expensive. Among the R packages created to run Bayesian linear models for genomic prediction, three main packages may be highlighted: BGLR-Bayesian Generalized Linear Regression (Pérez and de los Campos, 2014), BMTME-Bayesian Multi-Trait Multi-Environment (Montesinos-López et al., 2016) and BGGE-Bayesian Genotype plus Genotype by Environment (Granato et al., 2018). However, BGGE employs an optimization process that can be up to five times faster than BGLR and permits the incorporation more kernel structures than BMTME. For this reason, we implement the *kernel_model* function that runs the same optimization algorithm for Hierarchical Bayesian Modeling used in BGGE (see **Appendix 5**).

Below we describe a generic model structure that covers the diversity of possible combinations for modeling the phenotypic variation across MET. This model considers *k* genomic and *l* enviromic effects, plus fixed-effects and a random residual variation:

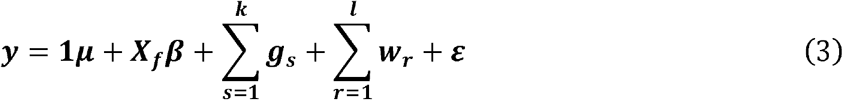

where ***y*** is the vector combining the means of each genotype across each one of the *q* environments in the experimental network, in which ***y*** = [***y***_1_, ***y***_2_,⋯ ***y_q_***]^***T***^. The scalar **1*μ*** is the common intercept or the overall mean. The matrix ***X_f_*** represents the design matrix associated with the vector of fixed effects ***β***. In some cases, this vector is associated with environmental effects (target as fixed-effect). Random vectors for genomic effects (***g_s_***) and enviromic-based effects (***w_r_***) are assumed to be independent of other random effects, such as residual variation (***ε***). It is a generalization for a reaction-norm model because, in some scenarios, the genomic effects may be divided as additive, dominance, and other sources (epistasis) and the genomic by environment (G×E) multiplicative effect. In addition, the envirotyping-informed data can be divided into several environmental kernels and a subsequent genomic by envirotyping (G×W) reaction-norm kernels. Based on Equation 6, the theory underpinning the *get_kernel* function is summarized in three

- **Benchmark Genotypic Effects**. These are baseline models accounting only for genotype-based effects, mostly associated with pedigree-based or genomic realized kinships. 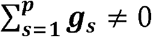 and 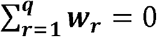, in which *g_s_* may be related to main genotype-effect (G) in the case of the main genotype-effect model (MM); and G plus a genotype by environment deviation (G+G×E), in the case of the so-called MDs model. Note that multiple genotype-relatedness kernels may be incorporated, e.g., for additive (A) and dominance (D) deviations and other sources of information from ‘omics.’ All genomic kernels must have the *p* × *p* dimension, in which *p* is the number of genotypes. However, this model does not consider any environmental effect.
- **Enviromic-enriched Main Effects**. We added the acronym ‘E’ to the MM and MDs models to denote ‘enviromic-enriched’ for EMM and EMDs models. These models consider 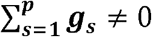 and 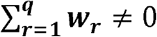, in which *g_s_* are related to G (EMM) or G+G×E (EMDs), and *w_r_* are only the main envirotype effects (W). In this type of model, the environmental effects can be modeled as a fixed deviation (using ***X_fβ_***) plus a random envirotyping-based variation 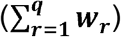).
- **Enviromic-based Reaction-Norm**. We added the acronym ‘RN’ for ‘reaction-norm’ to the MM and MDs models, resulting in RNMM and RNMDs models, respectively. As described in (ii), the environmental effects can now be modeled as fixed deviations (using ***X_f_β***) plus a random envirotyping-based variation 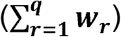. However, those RN models consider 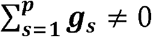 and 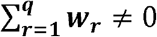, in which *g_s_* are related to G (RNMM) or G+G×E (RNMDs), and *w_r_* are related to main envirotype effects (W) plus an envirotype × genomic interaction (G×W). In this context, RNMM accounts for the variation due to G+W+GW, whereas RNMDS considers G+GE+W+GW.

#### Getting covariance structures with get_kernel

The *get_kernel* function has four main arguments: a list of genomic relationship kernels (**K_G**); a list of environmental relationship kernels (**K_E**); and a phenotypic MET data set (**data**), composed of a vector of environment identification (**env**), a vector of genotype identification (**gid**), and a vector of trait values (**y**); at last, the **model** argument sets the statistical model used (‘MM,’ ‘MDs’, ‘EMM’, ‘EMDs’, ‘RNMM’ and ‘RNMDs’). Each genomic kernel in **K_G** must have the dimension of *p* × *p* genotypes. This argument assumes K_G = NULL by default. If no structure for genetic effects is provided, the *get_kernel* function automatically assumes an identity matrix for genotype effects, in which it considers no relatedness among individuals. Finally, the argument **stage** (stage = NULL by default) states which development stages can be used to create *stage-specific enviromic kernels*. More detail about the latter is given in the Example 3 of the Results section.

In the same manner, **K_E** could have the dimension of *q* × *q* environments, but the environmental kernels can be built at the phenotypic observation level in some cases. It means that for each genotype in each environment, there is a different EC, according to particular phenology stages or envirotyping at the plant level. Thus, using the additional argument **dimension_KE** = c(‘q’, ‘n’) (default is ‘q’, for environment), the **K_E** may accomplish a kernel with *n* × *n*, in which *n* = *pq*. The basic usage of *get_kernel* is given below (**BOX 12**) and its detailed applications are provided in the Results section.

#### Modeling the phenotypic variation with kernel_models

Finally, the *kernel_model* function has four main arguments: a phenotypic MET data set (**data**), composed of a vector of environment identification (**env**), a vector of genotype identification (**gid**), and a vector of trait values (**y**), a list object for random effects (**random**, from *get_kernel*) and a matrix for fixed effects (**fixed**). For the Hierarchical Bayesian Modeling (see **Appendix 5**), the arguments for number of iterations (**iterations**, default is 1000) and the number of samples used for burn-in (**burnin**, default is 200) and thining (**thining**, default is 10) must be provided. The function has two main outputs: the predicted phenotypes (yHat), variance components for each random effect (VarComp). Below we show a brief example of the use of *kernel_model* for a MDs model (**BOX 13**) using the same inputs used in **BOX 12**.

#### BOX 12: Basic usage of get_kernel function

~~~
>data(“maizeYield”) # toy set of phenotype data (grain yield per environment)
>data(“maizeG”) # toy set of genomic relationship for additive effects
>data(“maizeWTH”) # toy set of environmental data
>y = “value” # name of the vector of phenotypes
>gid = “gid” # name of the vector of genotypes
>env = “env” # name of the vector of environments
>ECs = W_matrix(env.data = maizeWTH, var.id = c(“FRUE”,’PETP’,”SRAD”,”T2M_MAX”),statistic = ‘mean’)
## KG and KE might be a list of kernels
>KE = list(W = env_kernel(env.data = ECs)[[2]])
>KG = list(G=maizeG);
## Creating kernel models with get_kernel
>MM = get_kernel(K_G = KG, y = y,gid = gid,env = env, data = maizeYield,model = “MM”)
>MDs = get_kernel(K_G = KG, y = y,gid = gid,env = env, data = maizeYield, model = “MDs”)
>EMM = get_kernel(K_G = KG, K_E = KE, y = y,gid = gid,env = env, data = maizeYield, model = “EMM”)
>EMDs = get_kernel(K_G = KG, K_E = KE, y = y,gid = gid,env = env, data = maizeYield, model = “EMDs”)
>RMMM = get_kernel(K_G = KG, K_E = KE, y = y,gid = gid,env = env, data = maizeYield, model = “RNMM”)
>RNMDs = get_kernel(K_G = KG, K_E = KE, y = y,gid = gid,env = env, data = maizeYield, model = “RNMDs”)
~~~

#### BOX 13: Basic usage of kernel_model function

~~~
> fixed = model.matrix(~0+env, maizeYield)
> MDs = get_kernel(K_G = KG, y = y,gid = gid,env = env, data = maizeYield, model = “MDs”)
> fit = kernel_model(y = y,env = env,gid = gid, data = maizeYield,random = MDs,fixed = fixed)
~~~

### Practical Examples

Three practical examples were implemented to present a comprehensive overview of the most important functions of *EnvRtype*. First, we illustrate the use of *EnvRtype* for starting an envirotyping pipeline across different locations in the world (**Example 1**).

Second, we used the toy data set (*maizeG, maizeWTH* and *maizeYield*) to demonstrate different environmental similarities based on different environmental factors (**Example 2**). We used two envirotyping levels (per environment and per development stage at each environment) and two ECs (FRUE, PETP and FRUE+PETP) to demonstrate different ways to build environmental relatedness for GP. This type of application can be useful for researchers interested in predicting the individual genotypic responses shaped by genomic and enviromic-specific factors across existing experimental trials or for the assembly of virtual scenarios.

Finally, in **Example 3** we ran a genomic prediction study case in maize (*maizeG, maizeWTH* and *maizeYield*) involving three models (M1, Baseline Genomic MDs model; M2, Reaction Norm RNMM model and M3, Reaction Norm RNMM considering a different enviromic kinship for each development stage) and two cross-validation schemes (CV1: prediction of novel genotypes, using 20% of the data as a training set; CV00: prediction of novel genotypes at novel environments, using 3 of the 5 environments plus 20% of the genotypes as a training set).

## RESULTS

### Example1: Global-scale Envirotyping

To illustrate the use of *EnvRtype* for a global-scale envirotyping study, we consider different periods (and years) within the summer season at nine locations around the world (**BOX 14**): Goiânia (Brazil, 16.67 S, 49.25 W, from March 15th, 2020 to April 04th, 2020); Texcoco (Mexico, 19.25 N, 99.50 W, from May 15th, 2019 to June 15th, 2019); Brisbane (Australia, 27.47 S, 153.02 E, from September 15th, 2018 to October 04th, 2018); Montpellier (France, 43.61 N, 3.81 E, from June 18th, 2017 to July 18th, 2017); Los Baños (the Philippines, 14.170 N, 121.431 E, from May 18th, 2017 to June 18th, 2017); Porto-Novo (Benin, 6.294 N, 2.361 E, from July 18th, 2016 to August 18th, 2016), Cali (Colombia, 3.261 N, 76.312 W, from November 18th, 2017 to December 18th, 2017); Palmas (Brazil, 10.168 S, 48.331 W, from December 18th, 2017 to January 18th, 2018); and Davis (the United States, 38.321 N, 121.442 W, from July 18th, 2018 to August 18th, 2018). In this example, we use ‘GOI’, ‘TEX’, ‘BRI’, ‘MON’, ‘LOS’, ‘PON’, ‘CAL’, ‘PAL’ and ‘DAV’ to identify each location (**Figure 2A**).

**Figure 2.**
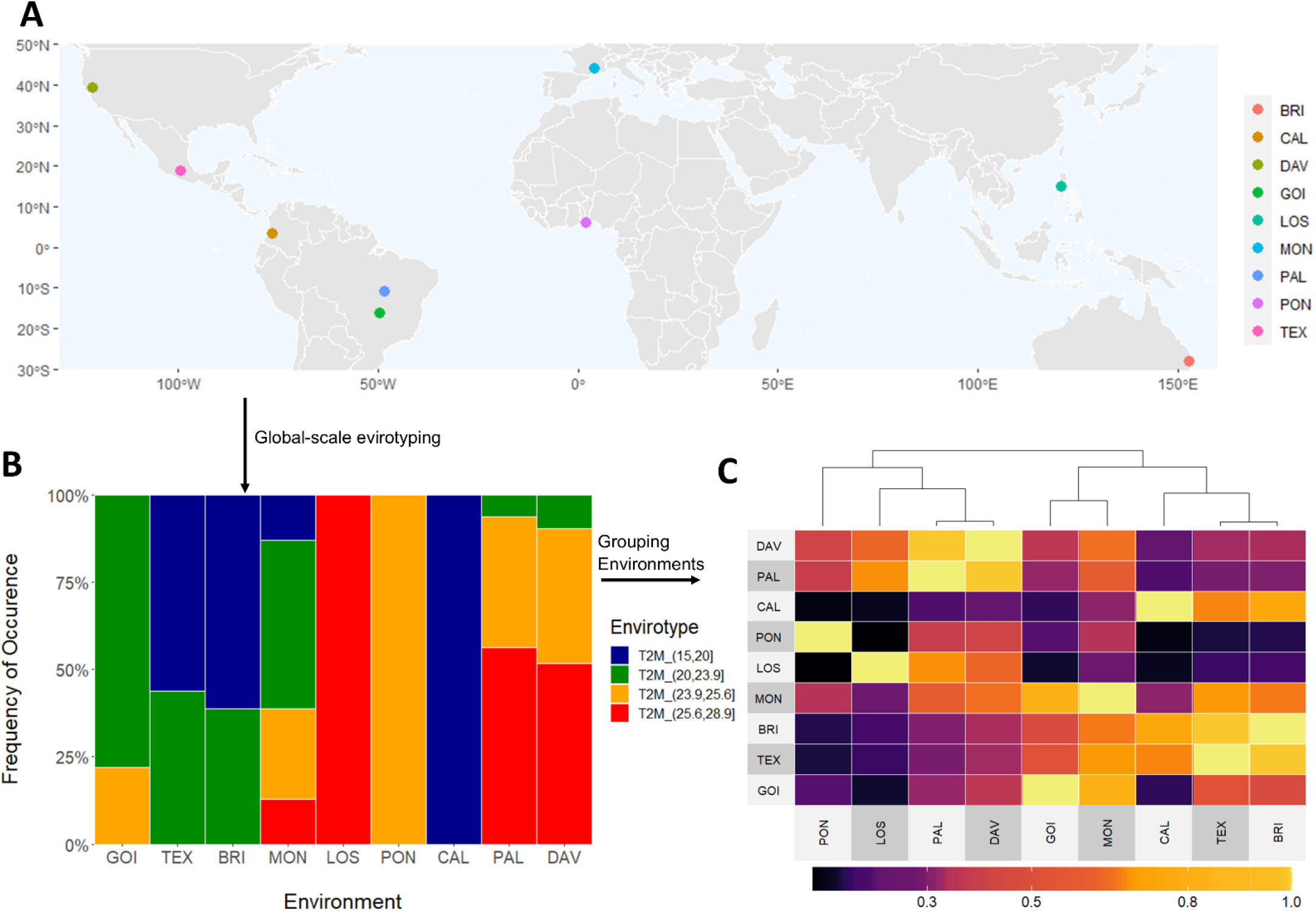
Workflow for a global-scale envirotyping analysis for air temperature effects in maize growing environments over diverse locations. **A.** Worldwide geographic positions of 9 locations used as toy examples. **B.** Panel of environmental types (ETs) for average air temperature during a specific month of a particular year in the summer season of each location. **C**. Environmental similarity matrix using Gaussian kernel method among the observed locations using the information of the observable ETs. From **A, B** and **C**, it is possible to highlight that environmental similarity among locations in different continents can be visualized to support the creation of a global-scale experimental network and support germplasm exchange between countries.

From the collected variables, it is possible to type any environmental factor or a core of environmental factors (**Figure 2B**). As a toy exemplification (**BOX 14–15**), we used the variable ‘T2M’ (the daily average temperature at 2 meters) to discover environmental types (ETs) and compute environmental similarity (**Figure 2C**). In this case, we used the Gaussian kernel as a sign of environmental distance, but it can also be used as kinship for predictive breeding (Costa-Neto et al., 2020a).

It is possible to see in this toy example, that perhaps locations in different continents might have similar ET trends for air temperature. This process can be done for several variables (single or joint) to better describe those similarities.

#### BOX 14: Remote Sensing for Several Places

~~~
> env = c(‘GOI’, ‘TEX’, ‘BRI’, ‘MON’, ‘LOS’, ‘PON’, ‘CAL’, ‘PAL’, ‘DAV’)
> lat = c(−16.67,19.25,-27.47,43.61,14.170,6.294,3.261,-10.168,38.321)
> lon = c(−49.25,-99.50,153.02,3.87,121.241,2.361,-76.312,-48.331,-121.442)
> start = c(‘2020-03-15’,’2019-05-15’,’2018-09-15’,
       ‘2017-06-18’,’2017-05-18’,’2016-07-18’,
       ‘2017-11-18’,’2017-12-18’,’2018-07-18’)
> end = c(‘2020-04-15’,’2019-06-15’,’2018-10-15’,
      ‘2017-07-18’,’2017-06-18’,’2016-08-18’,
      ‘2017-12-18’,’2018-01-18’,’2018-08-18’)
> env.data = get_weather(env.id = env, lat = lat, lon = lon, start.day = start, end.day = end)
~~~

#### BOX 15: Discovering ETs and similarity among locations

~~~
> ET = env_typing(env.data = env.data,env.id = ‘env’,var.id = ‘T2M’))
> EC = W_matrix(env.data = env.data,var.id = ‘T2M’)
> distances = env_kernel(env.data = ET,gaussian = T)[[2]]
> kinship = env_kernel(env.data = EC,gaussian = F, sd.tol = 3)[[2]]
~~~

### Example 2: Modeling Genomic-enabled Reaction-Norm

To illustrate the use of different ECs in modeling genomic-enabled reaction-norms, we ran a toy example involving a tropical maize data set available in *EnvRtype* (see **BOX 2**). From equation 3, we assumed the following baseline model: ***y*** = **1*μ*** + ***X_f_β*** + ***g*** + ***gE*** + ***ε***, where ***X_f_β*** is the fixed environmental effects, ***g*** is the random genomic-additive effects and ***gE*** is the genomic × environment interaction, modeled by a block diagonal matrix of genomic effects across environments (MDs model). Thus, we added enviromic effects following two envirotyping levels: (1) envirotyping mean values per environment (entire croplife), and (2) envirotyping for each time interval (development stage) across crop life, assuming fixed stages in terms of days after emergence. For each envirotyping level, we considered two types of ECs: the factor of temperature effect over radiation use efficiency (FRUE) and the difference between rainfall precipitation and crop evapotranspiration (PETP) (**Figure 3**). From equation (3), these matrices of ECs (EC1, EC2, EC3, EC4, EC5 and EC6) were arranged in three kernel structures using the RNMDs model, with the baseline genomic model updated to ***y*** = **1*μ*** + ***X_f_β*** + *g* + ***gE*** + ***EC*** + ***gEC*** + ***ε***, which resulted in 6 models (M1, M2, M3, M4, M5 and M6) according to each EC matrix used, plus a baseline genomic model (M0).

**Figure 3.**
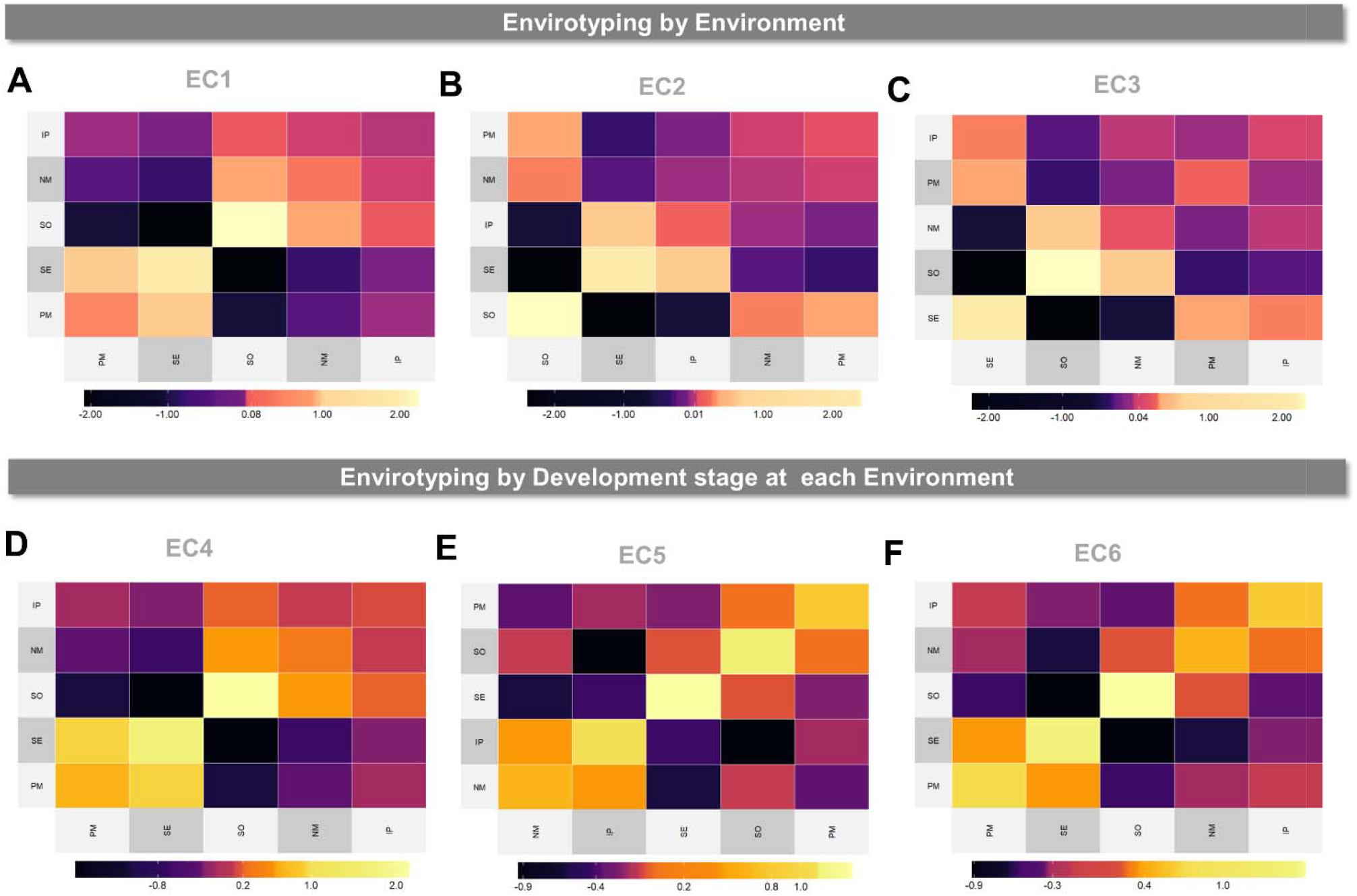
Linear enviromic kernels based on the combination of two environmental covariates (ECs) and two envirotyping levels (by environment and by development stage at each environment) for 5 locations (SE, PM, NV, IP and SO) across an experimental network of tropical maize. **A.** enviromic kernel considering only FRUE variable (impact of temperature on radiation use efficiency) at environmental level (EC1 matrix). **B.** enviromic kernel considering only the PETP variable (deficit of evapotranspiration, mm.day^-1^) for the entire crop life (EC2 matrix). **C.** enviromic kernel considering both FRUE and PETP for the entire crop live (envirotyping per environment, EC3 matrix). **D.** enviromic kernel considering only FRUE for each development stage at each environment (EC4 matrix). **E.** enviromic kernel considering only PETP for each development stage at each environment (EC5 matrix) and the combination of FRUE and PETP for each development stage (**D**, EC6 matrix).

**BOX 16** and **BOX 17** presents the codes to implement these envirotyping levels and model structures can be implemented in *EnvRtype*. Table 3 presents a brief summary of the variance components for each random effect. The inclusion of enviromic sources led to a drastic reduction of variance components for residual effects (from 0.837 in M0 to 0.262 in M6), and in some cases, the increase in the variance component for genomic effects (from 0.435 in M0 to 0.548 in M5). It was possible to observe that environmental variance is a key component that explains phenotypic variation. For the same raw environmental data, each envirotyping level and modeling structure impacts on modeling the phenotypic variation, such as comparing M1 to M4 and M2 to M5. Most of the variation due to G×E effects are better captured when some enviromic information is used in the model (from 0.329 in M0 to 0.786 in M3), leading us to infer that pure genomic G×E effects are inefficient in capturing the real pattern of genotype-environment differences observed in the phenotypic records. In the present example, the joint use of different EC led to the greatest reduction in error variance (M3 and M6), but the use of single ECs can be helpful in explaining genotype-specific difference as the G×E component is better estimated by considering the reaction-norm for particular environmental factors (e.g., FRUE in models M1 and M4). Finally, the effect of envirotyping level also impacted in the model’s ability in explaining phenotypic records, increasing the genomic variance (from 0.425 in M1 to 0.522 in M4, and from 0.387 in M2 to 0.548 in M5).

#### BOX 16: Envirotyping levels and model structures for modeling reaction-norm

~~~
> data(“maizeYield”) # toy set of phenotype data (grain yield per environment)
> data(“maizeG”) # toy set of genomic relationship for additive effects
> data(“maizeWTH”) # toy set of environmental data
> y = “value” # name of the vector of phenotypes
> gid = “gid” # name of the vector of genotypes
> env = “env” # name of the vector of environments
### 1-Environmental Covariables (ECs)
> stages = c(‘VE’, ‘V1_V6’, ‘V6_VT’, ‘VT_R1’, ‘R1_R3’, ‘R3_R6’, “H”)
> interval = c(0,7,30,65,70,84,105) # in days after emergence
> EC1 = W_matrix(env.data = maizeWTH, var.id = ‘FRUE’)
> EC2 = W_matrix(env.data = maizeWTH, var.id = ‘PETP’)
> EC3 = W_matrix(env.data = maizeWTH, var.id = c(‘FRUE✓PETP’))
> EC4 = W_matrix(env.data = maizeWTH, var.id = ‘FRUE’,
>   by.interval = T,time.window = interval,names.window = stages)
>EC5 = W_matrix(env.data = maizeWTH, var.id = ‘PETP’,
>   by.interval = T,time.window = interval,names.window = stages)### 2-Kernels
>K1 = list(FRUE = env_kernel(env.data = EC1)[[2]])
>K2 = list(PETP = env_kernel(env.data = EC2)[[2]])
>K3 = list(FRUE_PETP = env_kernel(env.data = EC3)[[2]])
>K4 = list(FRUE = env_kernel(env.data = EC4)[[2]])
>K5 = list(PETP = env_kernel(env.data = EC5)[[2]])
>K6 = list(FRUE_PETP = env_kernel(env.data = EC6)[[2]])
### 3-Obtain Kernel Models
>M0 = get_kernel(K_G = KG, y = y,gid = gid,env = env, data = maizeYield, model = “MDs”)
>M1 = get_kernel(K_G = KG, K_E = K1, y = y,gid = gid,env = env, data = maizeYield, model = “RNMDs”)
>M2 = get_kernel(K_G = KG, K_E = K2, y = y,gid = gid,env = env, data = maizeYield, model = “RNMDs”)
>M3 = get_kernel(K_G = KG, K_E = K3, y = y,gid = gid,env = env, data = maizeYield, model = “RNMDs”)
>M4 = get_kernel(K_G = KG, K_E = K4, y = y,gid = gid,env = env, data = maizeYield, model = “RNMDs”)
>M5 = get_kernel(K_G = KG, K_E = K5, y = y,gid = gid,env = env, data = maizeYield, model = “RNMDs”)
>M6 = get_kernel(K_G = KG, K_E = K6, y = y,gid = gid,env = env, data = maizeYield, model = “RNMDs”)
~~~

#### BOX 17: Running kernel_model to compute variance components

~~~
>fixed = model.matrix(~0+env, maizeWTH)
>iter = 1000; burn = 500; seed = 78172 # just as example
>model = paste0(‘M’, 0:6); Models = list(M0,M1,M2,M3,M4,M5,M6)
>Vcomp <- c()
>for(MODEL in 1:length(Models)){
>set.seed(seed)
> fit <- kernel_model(data = maizeYield,y = y,env = env,gid = gid,
>    random = Models[[MODEL]],fixed = Z_E,
>    iterations = iter,burnin = burn,thining = thin)
> Vcomp <- rbind(Vcomp,data.frame(fit$VarComp,Model = model[MODEL]))
>}
Vcomp
~~~

**Table 3.** Summary of variance components [and confidence intervals] for 7 genomic-based reaction-norm models with GxE (RNMDs), considering three envirotyping levels (no envirotyping, envirotyping by environment and envirotyping by development stage at environment) and three combinations of two environmental covariates (FRUE, PETP and FRUE+PETP). Models were fitted using all phenotypic records available (*n* = 150 genotypes at 5 environments = 750 records). Genomic kinships were based on additive effects. Enviromic kinships were built using a linear-covariance matrix (gaussian = FALSE). FRUE and PETP denote the covariates “effect of temperature on radiation use efficiency” (from 0 to 1) and the “difference between daily precipitation and daily evapotranspiration” (mm day^-1^), respectively.

| Envirotyping Level | Model | Random Effect |  |  |  |
| --- | --- | --- | --- | --- | --- |
|  |  | Environment (E) | Genotype (G) | GxE | Residual |
| No Envirotyping | M0 | - | 0.435<br>[0.397;0.480] | 0.329<br>[0.299;0.362] | 0.837<br>[0.763;0.924] |
| Envirotyping by environment | M1<br>(EC1 = FRUE) | 4.117<br>[3.789;4.493] | 0.425<br>[0.387;0.468] | 0.764<br>[0.696;0.843] | 0.849<br>[0.773;0.936] |
|  | M2<br>(EC 2 = PETP) | 3.440<br>[3.165;3.754] | 0.384<br>[0.350;0.423] | 0.786<br>[0.716;0.867] | 0.726<br>[0.662;0.801] |
|  | M3<br>(EC3 =FRUE+PETP) | 4.279<br>[3.938;4.670] | 0.497<br>[0.453;0.548] | 0.664<br>[0.605;0.733] | 0.456<br>[0.416;0.503] |
| Envirotyping by development stage at each environment | M4<br>(EC4 = FRUE) | 8.802<br>[8.099;9.605] | 0.522<br>[0.476;0.576] | 0.484<br>[0.441;0.534] | 0.266<br>[0.243;0.294] |
|  | M5<br>(EC5 = PETP) | 3.514<br>[3.233;3.835] | 0.548<br>[0.500;0.604] | 0.425<br>[0.388;0.469] | 0.267<br>[0.243;0.295] |
|  | M6<br>(EC6 = FRUE+PETP) | 1.595<br>[1.468;1.740] | 0.514<br>[0.468;0.566] | 0.464<br>[0.423;0.512] | 0.262<br>[0.238;0.289] |

### Example 3: Genomic Prediction using kernels for different development stages

Finally, we illustrate a case study of genomic prediction (GP) for two prediction scenarios (CV1 and CV00). In order to demonstrate the get_kernel function, we assumed a nonlinear kernel for enviromic effects (gaussian = TRUE). Different from the last example, in this study, we created different enviromic kernels for each development stage (M3). This model was compared to the benchmark reaction-norm model (M2, Jarquín et al., 2014) and the baseline multi-environment genomic model (M1, the MDs model López-Cruz et al., 2015). Thus, we ran the RNMM, which means that for M2 and M3 the genomic×environment effects are computed as Kronecker product between enviromic and genomic kinships, and for M1, as a block diagonal genomic matrix (MDs). In this example, we hope to demonstrate that for the same environment, it is possible to create different enviromic relatedness kernels according to the similarities found in different environments at the same development stage (**Figure 4**).

**Figure 4.**
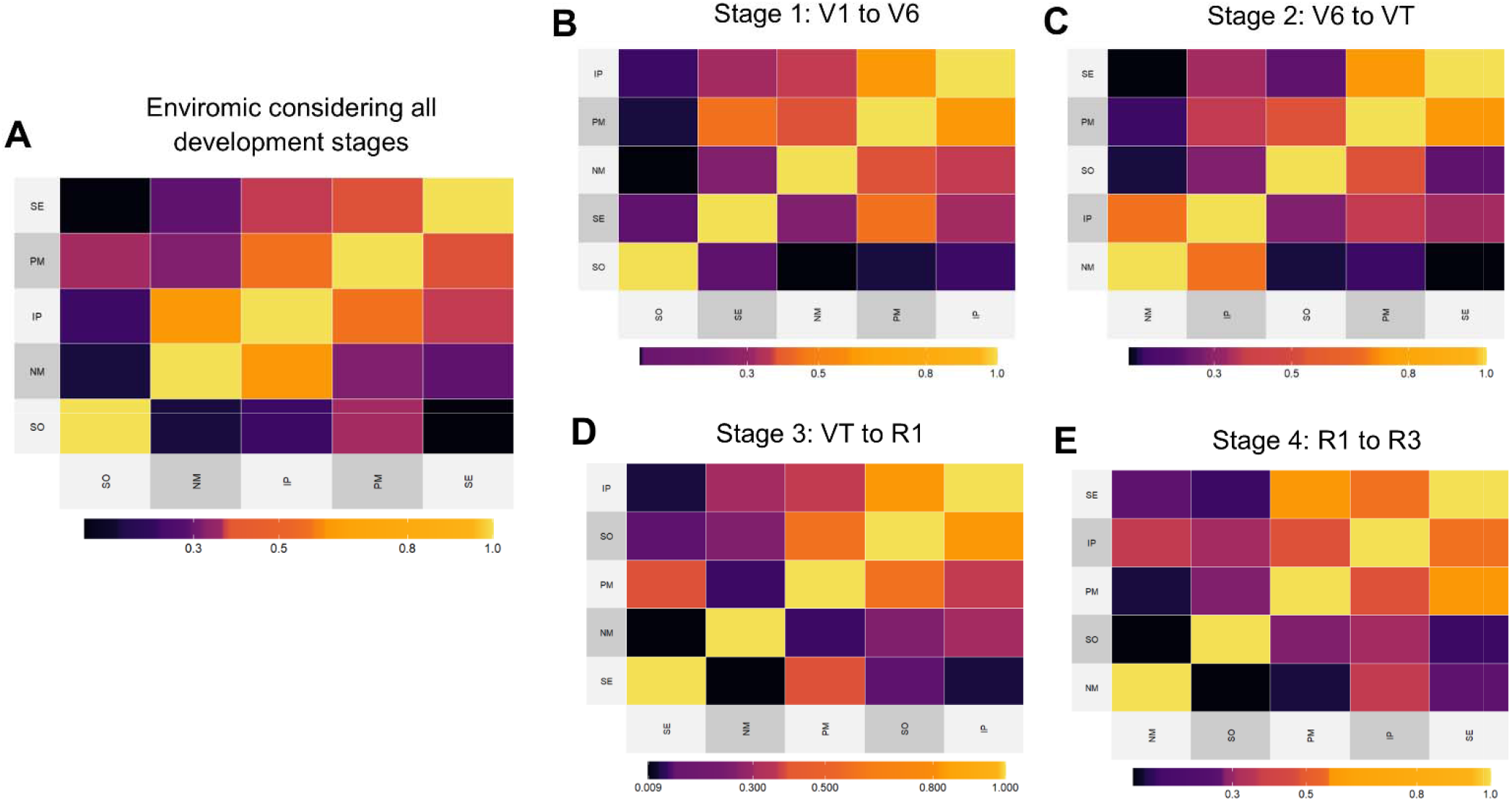
Nonlinear enviromic kernels (Gaussian) based on 13 environmental covariates over 5 tropical maize environments (locations (SE, PM, NV, IP and SO). **A.** enviromic kernel using a combination of 13 covariates at 4 development stages in maize (total of 91 ECs). **B.** enviromic kernel using variables 13 covariates at the initial vegetative stage (from V1 to V6). **C.** enviromic kernel using variables 13 covariates at the leaf growing stage (from V6 to VT). **D.** enviromic kernel using variables 13 covariates at the anthesis-silking interval (from VT to R1). **E.** enviromic kernel using variables 13 covariates at the grain filling interval (from R1 to R3).

We assume four development stages in maize: first vegetative stage (S_1_, from V1 to V6); second vegetative stage, in which the rate of biomass growth is increased in tropical maize (S_2_, from V6 to VT); anthesis-silking stage (S_3_, from VT to R1); and the grain filling stage (S_4_, from R1 to R3). To create this relatedness, 13 ECs were computed from daily weather data. The full matrix of ECs for all development stages considered a combination of 13 ECs by 4 development stages, thus resulting in 91 environmental descriptors (**BOX 18**).

#### BOX 18: Running kernel_model to compute variance components

~~~
## Organizing Environmental Covariables (ECs) in W matrix
> stages = c(‘VE’, ‘V1_V6’, ‘V6_VT’, ‘VT_R1’, ‘R1_R3’, ‘R3_R6’, “H”)
> interval = c(0,7,30,65,70,84,105); id.vars = names(maizeWTH)[c(10:15,23,25:30)] # variables used
>W.matrix = W_matrix(env.data = maizeWTH,env.id = ‘env’,
>    var.id = id.vars,by.interval = T,time.window = interval,
>    names.window = stages,center = F,scale = F)
## Kernel for the involving all development stages
>K_F <- env_kernel(env.data = W.matrix,gaussian = T)[[2]]
## Kernels for each development stage
>K_S <- env_kernel(env.data = W.matrix,gaussian = T,stages = stages[2:5])[[2]]
# K_G (genotype) and K_E (environment) must be a list of kernels
>K_G = list(G = maizeG); K_F <- list(E = K_F)
## Assembly Genomic and Enviromic Kernel Models
>M1 = get_kernel(K_G = K_G,data = maizeYield,env = env,gid = gid,y = y, model = “MDs”) # baseline model
>M2 = get_kernel(K_G = K_G, K_E = K_F, data = maizeYield,env = env,gid = gid,
>    y = y, model = “RNMM”,dimension_KE = ‘q’) # reaction-norm 1
>M3 = get_kernel(K_G = K_G, K_E = K_S, data = maizeYield,env = env,gid = gid,
>    y = y,model = “RNMM”,dimension_KE = ‘q’) # reaction-norm 2
~~~

Figure 3 shows that for the same five environments, the crops’ growing conditions, and consequently the patterns of similarity, differ according to the crop development stage. So, it’s feasible to hypothesize a relatedness build up for some development stages may be more informative of the environmental-phenotype covariance among field trials than others, and perhaps more so than all environmental variables at all development stages. The biological explanation behind this hypothesis relies on ecophysiology, in which the plants are more or less sensitive to environmental variations at specific development stages. If the growing conditions drastically differ at some key development stage and do not differ in others that do not have a strong impact on the final phenotypic expression, it is feasible to assume that the environmental variance for those ‘non environmental polymorphic stages may lead to an increased noise in the enviromic relatedness, reducing then the accuracy of GP using reaction-norms. In order to test this hypothesis, we ran the cross-validation (**BOX 19**) for two prediction scenarios (CV1 and CV00). Below we present an example of code for running GP. More detail about the codes is given in the Supplementary Codes section.

#### BOX 19: Genomic Prediction using kernel_model

~~~
>source(‘https://raw.githubusercontent.com/gcostaneto/SelectivePhenotyping/master/cvrandom.R’)
>rep = 10; seed = 1010; f = 0.20; iter = 5E3; burn = 1E3; thin = 10; Y = maizeYield
>TS = Sampling.CV1(gids = Y$gid,f = f,seed = seed,rep = rep,gidlevel = F)
>fixed = model.matrix(~0+env,Y)
> require(foreach); require(EnvRtype)
>results <-foreach(REP = 1:rep, .combine = “rbind”)%:%
>foreach(MODEL = 1:length(model), .combine = “rbind”)%dopar% {
>yNA <-Y
>tr <-TS[[REP]]
>yNA$value[-tr] <- NA
>fit <- kernel_model(data = yNA,y = y,env = env,gid = gid,
>     random = Models[[MODEL]],fixed = fixed,
>     iterations = iter,burnin = burn,thining = thin)
> df<-data.frame(Model = model[MODEL],rep=REP,
>  rTr=cor(Y$value[tr], fit$yHat[tr],use = ‘complete.obs’),
>     rTs=cor(Y$value[-tr], fit$yHat[-tr],use = ‘complete.obs’))
> return(df)
>}
~~~

Table 4 presents a summary of the variance components for the random effects of each model. An increased trend in the genomic variance was observed as the inclusion of some enviromic data (from 0.426 in M1 to 0.509 in M2 and 0.555 in M3) and reduction of residual error variance (from 0.848 in M1 to 0.269 in M2 and 0.262 in M3). Despite that, for this germplasm evaluated at this experimental network, the effect of G×E decreased when estimated using a enviromic kinship (from 0.353 in M1 to 0.269 in M2), but were better understood when dissected for each development stage (M3). The variance components for environmental relatedness plus genomic×stage interaction was higher for reproductive-related stages (S_3_ and S_4_), suggesting that these stages may be more important in explaining environmental-phenotype covariances across field trials than using all environmental information to build up enviromic kinships.

**Table 4.**
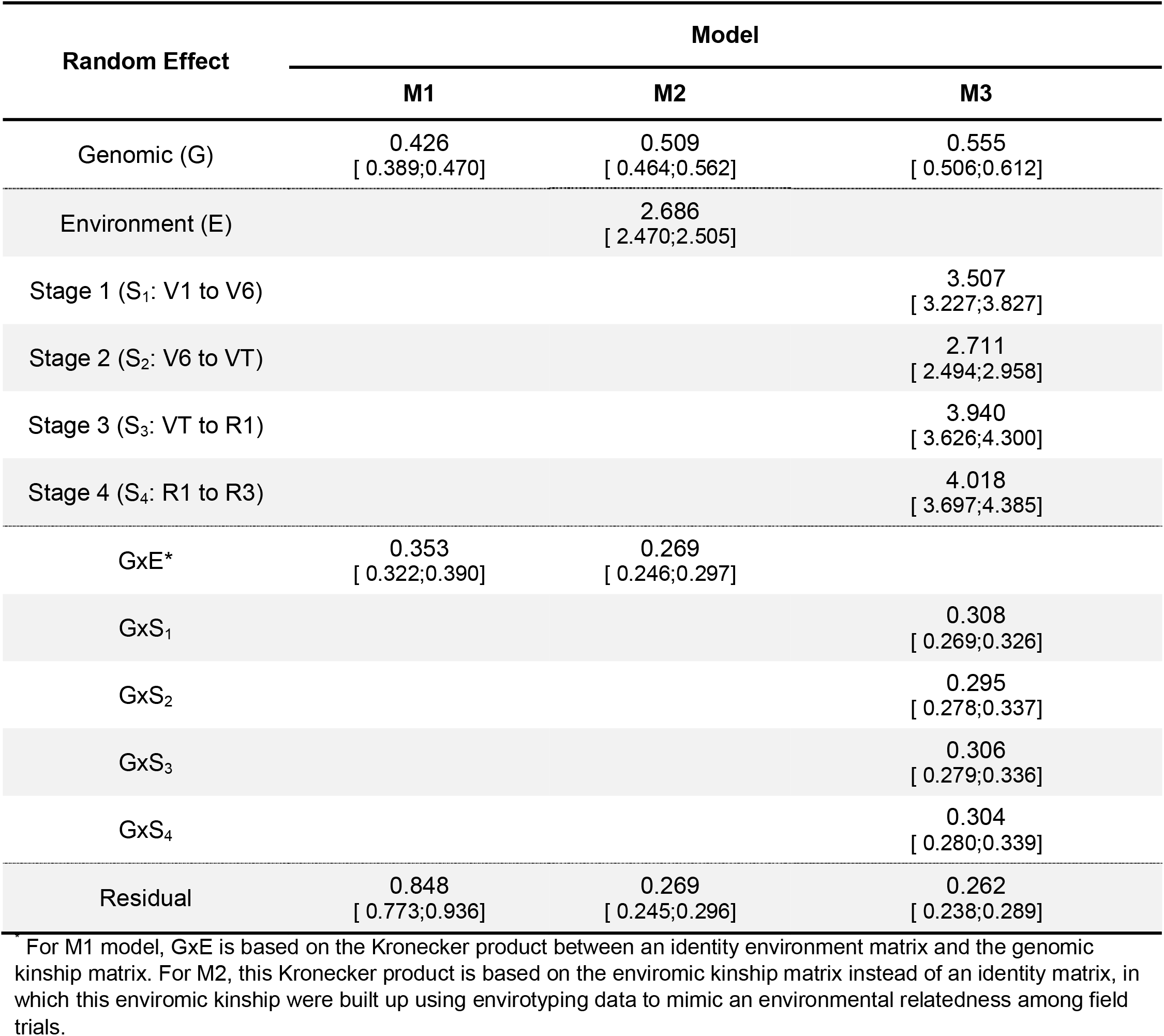
Summary of the variance components for three modeling structures (M1, Baseline Genomic Model; M2, Benchmark Reaction-Norm model; M3, Reaction-Norm for each development stage) considering different sources of phenotypic variation due to genomic and enviromic effects. Confidence intervals (α = 5%) for each variance component is given between square brackets. Horizontal dashed lines separate the genomics, environmental and genomic × environment effects. Genomic kinships were based on additive effects. Enviromic kinships were built using a nonlinear method (gaussian = TRUE).

| Random Effect | Model |  |  |
| --- | --- | --- | --- |
|  | M1 | M2 | M3 |
| Genomic (G) | 0.426<br>[ 0.389;0.470] | 0.509<br>[ 0.464;0.562] | 0.555<br>[ 0.506;0.612] |
| Environment (E) |  | 2.686<br>[ 2.470;2.505] |  |
| Stage 1 (S<sub>1</sub>: V1 to V6) |  |  | 3.507<br>[ 3.227;3.827] |
| Stage 2 (S<sub>2</sub>: V6 to VT) |  |  | 2.711<br>[ 2.494;2.958] |
| Stage 3 (S<sub>3</sub>: VT to R1) |  |  | 3.940<br>[ 3.626;4.300] |
| Stage 4 (S<sub>4</sub>: R1 to R3) |  |  | 4.018<br>[ 3.697;4.385] |
| GxE* | 0.353<br>[ 0.322;0.390] | 0.269<br>[ 0.246;0.297] |  |
| GxS<sub>1</sub> |  |  | 0.308<br>[ 0.269;0.326] |
| GxS<sub>2</sub> |  |  | 0.295<br>[ 0.278;0.337] |
| GxS<sub>3</sub> |  |  | 0.306<br>[ 0.279;0.336] |
| GxS<sub>4</sub> |  |  | 0.304<br>[ 0.280;0.339] |
| Residual | 0.848<br>[ 0.773;0.936] | 0.269<br>[ 0.245;0.296] | 0.262<br>[ 0.238;0.289] |

\* For M1 model, GxE is based on the Kronecker product between an identity environment matrix and the genomic kinship matrix. For M2, this Kronecker product is based on the enviromic kinship matrix instead of an identity matrix, in which this enviromic kinship were built up using envirotyping data to mimic an environmental relatedness among field trials.

Finally, in Table 5, we present the accuracy of the statistical models for each prediction scenario. These results were obtained by the average Pearson’s Moment Correlation (*r*) for each one of the 30 random samples of training sets. The use of enviromic information was beneficial for both prediction scenarios. The ability to predict novel genotypes at known growing conditions (CV1), using only the phenotypic records of 20% of the germplasm led to an increase from *r* = 0.130 (baseline genomic model, M1) to *r* = 0.762 (benchmark reaction-norm model, M2), in which the latter was not different for the reaction-norm accounting for stage-specific enviromic kernels (*r* = 0.760 for M3). However, great differences between M2 and M3 were observed for predicting novel genotypes at novel growing conditions (CV00). In this scenario, based on the phenotypic records from 20% of the germplasm evaluated in 3 of the 5 environments (so the remaining 2 were used as testing-environments), the M3 model outperformed all models (*r* = 0.504) in comparison to M2 (*r* = 0.485) and M1 (*r* = 0.102). This last model has the worst performance due to the lack of phenotypic records.

**Table 5.** Accuracy (± standard deviation) of statistical models for phenotype prediction using genomic (M1) and genomic plus enviromic sources of variation (M2 and M3) in predicting novel genotypes at known environments (CV1), using 20% of the genotypes as a training set, and novel genotypes and novel environments (CV00), used as a training set 20% of the genotypes phenotyped at 3 from the 5 environments.

| Model | Prediction Scenario |  |
| --- | --- | --- |
|  | CV1 | CV00 |
| M1 (Baseline Genomic × Environment) | 0.130 ± 0.047 | 0.102 ± 0.045 |
| M2 (Benchmark Reaction-Norm) | 0.762 ± 0.024 | 0.485 ± 0.211 |
| M3 (Reaction-Norm for Each Development Stage) | 0.760 ± 0.028 | 0.504 ± 0.194 |

## DISCUSSION

The collection, processing, and use of envirotyping data in genomic-based studies do not depend only on the quality of the data sources. Here we demonstrate that the increased ecophysiological knowledge in envirotyping increases statistical models’ accuracy in genomic prediction and provides a better explanation of the sources of variation, while increasing those efficiency models. The correct use of envirotyping data depends on the quality of data processing and is specific for each crop species (or living organism). The same ‘environment’ (considering a time interval for a target location) may result in different environmental types (ETs) for each organism, depending on their sensitivity to constant and transitory environmental variations. Thus, in this study, we presented some of those concepts and created functions (and gathered others from different R packages) to facilitate the use of envirotyping data in quantitative genomics.

We presented a user-friendly software, but also a cost-effective pipeline aimed at democratizing the use of envirotyping in several fields of plant research. *EnvRtype* is an opensource package and all advances made up until now are freely available, a situation that can ultimately boost the predictive breeding for low-budget research programs to invest in environmental sensors or perform experiments across a geographically heterogeneity region. Thus, as the remote sensing tools and databases evolve, the power of *EnvRtype* to perfom a quick and accurate envirotyping pipeline also evolves. In addition, other types of data sources can easily be integrated in the modeling approaches. For example, the use of high-throughput phenotypes can easy be integrated in predictive models by using *get_kernel* to build phenomics-realized relationship kernels (Rincent et al., 2018; Crain et al., 2018; Cuevas et al., 2019). Other kernel methods, such as Deep Kernel (Cuevas et al., 2019; Costa-Neto et al., 2020a), can be used to create kernels to be incorporated in Bayesian kernel models using *kernel_model* function. Thus, despite the provided end-to-end pipeline to interplay envirotyping in quantitative genomics, the users can adapt their codes and integrate different sources of information using the *EnvRtype* functions, such other omics data (Westhues et al., 2017).

We also showed that global envirotyping networks could be built using remote sensing tools and functions provided in *EnvRtype*. The combination of remote sensing + typing strategies is a powerful tool for turbocharging global partnerships of field testing and germplasm exchange. It also contributes to increasing the prediction of genotypes across a wide range of growing conditions, i.e., the so-called adaptation landscapes (Messina et al., 2018; Bustos-Korts et al., 2019). This can involve past trends and virtual scenarios (Gillberg et al., 2019; de los Campos et al., 2020). Associated with predictive GIS tools, the recommendation of cultivars could also be leveraged for specific regions (Costa-Neto et al., 2020b). It could also increase a better definition of field trial positioning (Rincent et al., 2017; Resende et al., 2020) and how breeding strategies have impacted crop adaptation in the past (Heinemann et al., 2015. Evidence of this suggestion is the increased ability of predicting novel genotypes at novel growing conditions achieved by obtaining a deeper understanding of how environments are more or less related at each development stage of crop life.

Despite the benefits and potential uses of *EnvRtype*, we can envisage the following limitations: (1) the resolution of satellite-based weather system (Nasa Power data base), corresponding to 0.5°×0.5° (~55,5 km × 55,5 km) of longitude by latitude, may compromise the discrimination of environments in close geographical proximity; (2) the quality of point-estimates of environmental data using *extract_GIS* function from public GIS databases depends on the file resolution available; (3) the need for a good registration of geographic coordinates of the target environment, but also on knowing the ‘window’ between harvest and sowing (for agricultural crops); (4) management factors must be included manually in *W_matrix*, which we strongly suggest in order do avoid mistakes; (5).

## APPENDIX

## Appendix 1. What is ‘environment’ and how to remotely collect environmental data

We use the term ‘environment’ to refer to delimited unit that combines location, planting date, and management, which gathers the fluctuation for a core of environmental factors. Thus, the first step of any envirotyping study is to collect reliable environmental data. However, for most breeding programs worldwide, this step is limited by the availability of sensing equipment (e.g., weather stations) installed in the field or nearby site. It is important to highlight that some equipment can be expensive or difficult to access for some research groups in certain regions, more specifically developing countries. For this reason, below, we present two justifications for incorporating a remote environmental sensing routine (in-silico) into this package. Then, we present recommendations to enrich the envirotyping platforms to collect and organize environmental data that will be useful to breeders’ decision making.

Firstly, in order to facilitate the step of collecting environmental data, we decided to include a routine for collecting raw daily weather data through the NASA POWER database (https://power.larc.nasa.gov/), which can access daily information anywhere on earth. This database was integrated using the tools provided by the nasapower R package (Sparks, 2018). Additionally, we integrated the raster R package to support downloading climatic data (from the WorldClim database) and SRTM (Shuttle Radar Topography Mission, which provides elevation information). The information from both databases is freely available and can be downloaded using geographical coordinates (latitude and longitude, given in decimal degrees, both in WGS84 format) for a specific window of time (e.g., from sowing to harvest).

Secondly, the collected environmental data processing requires some expertise in fields such as agrometeorology, soil physics, and ecophysiology. For this to be useful in explaining crop adaptation, the environmental data must be representative of some envirotype-to-phenotype dynamic linked to certain ecophysiological knowledge (e.g., air temperature, relative air humidity, and solar radiation driving the crops’ evapotranspiration and, consequently, the soil-water balance). A direct example of the importance of processing raw envirotyping data into ecophysiological enriched information is given for the ‘daily air temperature’ variable. This variable can be processed in heat units, heat stress effects on radiation use efficiency, and thermal range, which is species-specific. For some traits such as grain yield in maize, the impact of temperature-derived factors differs from what is observed in traits, such as plant height or flowering time. This dynamic also varies across crop developmental stages, which can be more or less prone to become a stress factor at certain stages (e.g., in maize, heat at the flowering time has a more significant impact on grain yield). Below is a detailed description of some of those subroutines.

## Appendix 2. Concepts underlying Radiation effects in Plants

The radiation balance in crop systems is regulated by the difference between the amount of incident radiation, absorbed energy by the plants and soil surface, and the converted thermal energy. From Nasa Power, the radiation outputs are given in terms of Top-of-atmosphere Insolation (ALLSKY_TOA_SW_DWN), Insolation Incident on a Horizontal Surface (Shortwave, ALLSKY_SFC_SW_DWN), and Downward Thermal Infrared Radiative Flux (Longwave, ALLSKY_SFC_LW_DW). Thus, the net solar radiation available for the physiological process of growth (biomass production) is given by the difference between longwave and shortwave, i.e., *SRAD* = ALLSKY_SFC_LW_DW – ALLSKY_SFC_SW_DWN, in MJ m^-2^ d^-1^. It is possible to download more solar-related parameters directly from Nasa Power website, (https://power.larc.nasa.gov/data-access-viewer/).

In most growth modeling approaches, the effect of radiation use efficiency (RUE) is the main target to describe the relationship between the available energy in the environment and how the plants translate it into biomass (see subsection about thermal parameters). In this context, this environmental variation source is important to understand the differences in potential yield observed in genotypes evaluated across diverse environments. Radiation is also vital as a source for regulating the available energy for other biophysical processes, such as evaporation, transpiration, and temperature (see subsection Processing Atmospheric Parameters).

## Appendix 3. Concepts underlying the Effect of Temperature in Plants

Thermal variables are essential for regulating the rates of critical biochemical processes within an organism. At the cell level, the effect of temperature may regulate the rate of enzymatic reactions, in which critical values may lead to denaturation of those enzymes and the death of the cell. At the plant level, temperature-related variables regulate the balance between photosynthesis (gross and net) and respiration in the canopy, impacting radiation use efficiency (RUE). It is also related to the transpiration rates and, consequently, to the absorption of nutrients from water flux in the roots. At the reproductive stages, temperature affects the efficiency of pollination, which is directly related to the crop’s final yield, especially for species in which grain yield is the main target trait. Phenology development rates are also strongly influenced by temperature (e.g., growing degree-days, *GDD*), in which the balance between biomass accumulation and acceleration of the crop cycle may compromise the source: sink relations and then the final yield.

Table 2 summarizes the cardinal limits of temperature for several species. Those cardinal limits are used to compute growing degree-days (*GDD*) and the effect of temperature on radiation use efficiency (*FRUE*). The first is useful to predict phenological development, while the second is an ecophysiology parameter used to quantify the impact of temperature on crop growth and biomass accumulation in crop models (Soltani and Sinclar, 2012). Thus, both can be useful to relate how temperature variations shape some species’ adaptation in the target environment. *GDD* is also important for modeling plant-pathogen interactions because some pests and diseases have temperature-regulated growth.

In this context, the dew point (*T2MDEW*) is another agrometeorological factor that is greatly important for crop health. This factor determines the establishment of diseases (especially fungus) under the leaf to being related to the evaporation process in the stomata. Finally, the daily temperature range (*T2M_RANGE*) impacts processes such as floral abortion in crops where the main traits are related to grain production. For more details about the impact of temperature on diverse crops, please check Luo (2011).

## Appendix 4. Concepts underlying the Effect of Atmospheric demands in Plants

The dynamics of precipitation (rainfall) and water demand (evaporation+plant transpiration) are regulated as a consequence of the balance of radiation and thermal-related processes in the atmosphere (Soltani and Sinclair, 2012; Allen et al., 1998). The soil-plant-atmosphere continuum involves water dynamics from the soil, passing through plant tissues, and going back to the atmosphere through the stomata. This process’s rate is deeply related to the biomass production of plants and the absorption of nutrients by the mass flux in roots. Because of that, water demands are essential for measuring the quality of some growing environments.

Here we used the Priestley-Taylor equation to compute the reference crop evapotranspiration. With this equation, the empirical constant (alpha = α) may range from 1 (at humidity conditions) to 2 (at arid conditions). First, we compute the vapor pressure, determined by *e_a_* = *RH* × *e_s_* (Dingman, 2002), where *e_s_* is the saturation vapor pressure defined as (Buck, 1981):

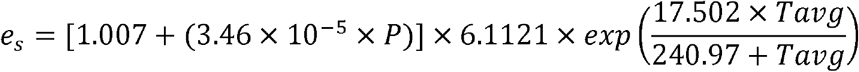

where *Tavg* is the average air temperature, and *P* is the air pressure (kPa) computed from elevation as *P* = 101.3 x (293 – 0.0065 x *ALT*/293)^5.26^. Thus, from the daily vapor pressure (*e_a_*), we compute the slope of the saturation vapor pressure curve (Δ), by (Dingman, 2002):

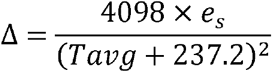

Finally, the reference evapotranspiration (ET_0_) is computed as:

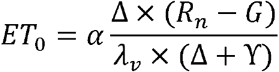

where *λ_v_* is the volumetric latent heat of vaporization (2453 MJ m^-3^) and □ is the psychometric constant (kPa C^-1^), that can be computed from air pressure as γ = 0.665 x 10^−3^*P* (Allen et al., 1998). For crops, we encourage the use of crop coefficients (K_c_, dimensionless) to translate *ET*_0_ in crop-specific evapotranspiration. This K_c_ is computed from empirical phenotypic records (crop height, the albedo of the soil-crop surface, canopy resistance) combined with in-field sensors (evaporation from the soil) or using K_c_ estimates for each crop species. Allen et al. (1998) provide a wide number of general K_c_ values to be used in this sense. For a complete understanding of soil-water dynamics, we suggest using pedotransfer functions to derive some hydraulic properties of the soil, such as infiltration rate and water retention parameters. It can be done by soil samples or from remotely collected data from SoilGrids using *extract_GIS()*.

## Appendix 5. Hierarchical Bayesian Modeling used in kernel_model

In this appendix, we present the hierarchical Bayesian modeling used in *kernel_model* function of *EnvRtype*. From the package for Bayesian Genotype plus Genotype × Environment (*BGGE*), which contains a function called *BGGE*(), we collected the main code and adapted it for our purposes. This function aims to solve mixed linear models through Hierarchical Bayesian Modeling--more detail about that can be found at Granato et al. (2018). Thus, we integrated the packages *EnvRtype* with *BGGE* into a single platform. If the users want to run genome-enabled models without enviromic data, we strongly suggest the use of *BGGE*() instead of *kernel_models*() because the *BGGE* package permits the construction of other modeling structures beyond the MM and MDs presented in this study. Below, we briefly describe the main distributions and priors used by this package.

The algorithm starts with a reparameterization of each variance-covariance matrix (***K***) provided by using the *get_kernel*() function. Each ***K*** is reparametrized using an eigen-decomposition procedure as suggested by De Los Campos et al. (2010), in which ***K*** = ***USU***’, where ***S*** is a diagonal matrix with n non-zero eigenvalues and ***U*** is an orthogonal matrix with eigenvectors. Then, an orthogonal transformation is applied to increase the computational efficiency of the further steps of the Bayesian approach. This transformations consists of a phenotypic parametrization, represented as ***d*** = ***U***’***y***, and any kernel-based random effect (***b*** = ***U***’***u***) and error variation (**e** = ***U***′***ε***) is now represented into a reparametrized normal distribution as 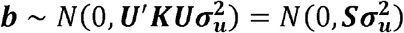 and 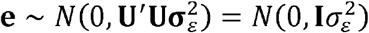 (Cuevas et al., 2017, 2019). Finally, the distribution of the transformed data is given by:

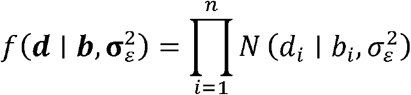

where the acronym *i* now denotes each random effect (variance-covariance) considered (from *get_kernel*). As this Bayesian linear model assumes 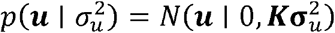, the conditional of any *b_i_* is given as 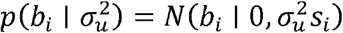, where *s_i_* are the eigenvalues.

Thus, we assume a conjugate prior distribution of 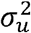 and 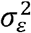, given by inverse chi-squared with, 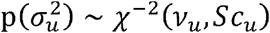 and 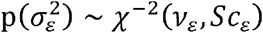 respectively, in which *v_u_* and *v_ε_* denote the degree of freedom, and *Sc_u_* and *Sc_ε_* the scale factors for ***u*** and ***e***. Finally, The Markov Chain Monte Carlo (MCMC) procedure is then used to generate the conditional distributions through a Gibbs sampler using the joint posterior distribution 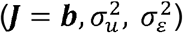, given the parameters (***P*** = ***d***, *v_u_, v_ε_, Sc_u_, Sc_ε_* and ***S***) as:

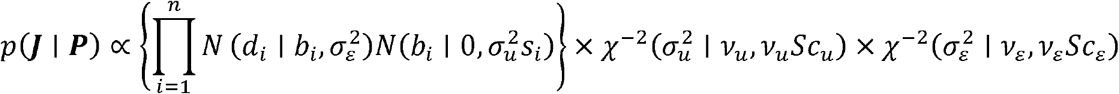

